# Proteostasis and metabolic dysfunction in a distinct subset of storage-induced senescent erythrocytes targeted for clearance

**DOI:** 10.1101/2024.09.11.612195

**Authors:** Sandy Peltier, Mickaël Marin, Monika Dzieciatkowska, Michaël Dussiot, Micaela Kalani Roy, Johanna Bruce, Louise Leblanc, Youcef Hadjou, Sonia Georgeault, Aurélie Fricot, Camille Roussel, Daniel Stephenson, Madeleine Casimir, Abdoulaye Sissoko, François Paye, Safi Dokmak, Papa Alioune Ndour, Philippe Roingeard, Emilie-Fleur Gautier, Steven L Spitalnik, Olivier Hermine, Pierre A Buffet, Angelo D’Alessandro, Pascal Amireault

## Abstract

Although refrigerated storage slows the metabolism of volunteer donor RBCs, cellular aging still occurs throughout this *in vitro* process, which is essential in transfusion medicine. Storage-induced microerythrocytes (SMEs) are morphologically-altered senescent RBCs that accumulate during storage and which are cleared from circulation following transfusion. However, the molecular and cellular alterations that trigger clearance of this RBC subset remain to be identified. Using a staining protocol that sorts long-stored SMEs (i.e., CFSE^high^) and morphologically-normal RBCs (CFSE^low^), these *in vitro* aged cells were characterized.

Metabolomics analysis identified depletion of energy, lipid-repair, and antioxidant metabolites in CFSE^high^ RBCs. By redox proteomics, irreversible protein oxidation primarily affected CFSE^high^ RBCs. By proteomics, 96 proteins, mostly in the proteostasis family, had relocated to CFSE^high^ RBC membranes. CFSE^high^ RBCs exhibited decreased proteasome activity and deformability; increased phosphatidylserine exposure, osmotic fragility, and endothelial cell adherence; and were cleared from the circulation during human spleen *ex vivo* perfusion. Conversely, molecular, cellular, and circulatory properties of long-stored CFSE^low^ RBCs resembled those of short-stored RBCs.

CFSE^high^ RBCs are morphologically and metabolically altered, have irreversibly oxidized and membrane-relocated proteins, and exhibit decreased proteasome activity. *In vitro* aging during storage selectively alters metabolism and proteostasis in SMEs, targeting these senescent cells for clearance.

## Introduction

Transfusion medicine affects millions of patients worldwide, with ∼119 million blood collections annually. In most European countries, whole blood donations are processed into RBC concentrates, and stored at 4°C in saline-adenine-glucose-mannitol (SAGM) solution for up to 42 days. Despite slowing of their metabolism during refrigerated storage, RBCs continue to age *in vitro*. This time-related decline share similarities with physiological senescence *in vivo*. One important difference between these two aging processes is that storage-induced senescent RBCs accumulate alterations in the confined and protective environment of the storage bag, thereby (temporarily) avoiding clearance from the circulation; in contrast, newly-senescent RBCs *in vivo* are continuously cleared from circulation. Multiple storage-related RBCs modifications have been described, reflecting the deterioration in their quality (1, 2). As examples, energy metabolism progressively diminishes (3–6), with decreased intracellular levels of ATP (7–9) and antioxidants (10), followed by increasing oxidant stress (2, 11). Thus, refrigerator-stored RBCs are less able to cope with oxidative stresses generated during storage, leading to increased lipid and protein oxidation (5, 12–17). These metabolic and oxidative stresses progressively modify multiple RBC properties, including increased surface exposure of phosphatidylserine (PS) and endothelial cell adherence, decreased osmotic resistance and deformability, and altered morphology (18–21). The evolution of storage lesion markers in individual RBC units during *in vitro* aging varies greatly, with donor sex, age, and genetics influencing end-of-storage oxidative and spontaneous hemolysis *in vitro* and post-transfusion RBC recovery and hemoglobin increments *in vivo* (22–25).

Although recent prospective studies did not show a survival advantage when transfusing short-stored RBC concentrates (*vs* standard of practice) (26–30), the storage lesion remains a matter of concern as it is responsible for the rapid clearance of a significant proportion of transfused RBCs, thereby decreasing transfusion efficacy (25, 31–35). Nonetheless, these studies suggest that only a fraction of the stored RBCs is sufficiently altered *in vitro* to be recognized, and then cleared, post-transfusion *in vivo*, by the recipient’s mononuclear phagocyte system.

Decreased intracellular ATP (36–39), one component of the RBC storage lesion, negatively correlates with transfusion recovery, indicating that *in vitro* markers can inform on post-transfusion outcomes. However, this metabolite was measured at the whole-population level in RBC units; thus, its evolution in individual RBCs is unknown. Similarly, storage-induced morphological alterations are a cellular marker of post-transfusion clearance. Indeed, storage-induced micro-erythrocytes (SMEs; comprising type III echinocytes, sphero-echinocytes, and spherocytes) accumulate during storage, vary between donors, negatively correlate with transfusion recovery in healthy volunteers, and are preferentially cleared from the circulation *in vivo* in a mouse transfusion model (40). However, the molecular and cellular abnormalities occurring in this distinct subset of morphologically-altered RBCs, triggering their post-transfusion clearance, remain to be identified. Using a simple CFSE staining protocol (41) combined with flow cytometric sorting, preparations were obtained containing either >90% SMEs (i.e., CFSE^high^) or >95% long-stored morphologically-normal cells (i.e., CFSE^low^), thereby allowing in-depth comparisons of their properties.

The current study characterizes SMEs, and compares them with morphologically-normal long-stored and short-stored RBCs. To this end, molecular phenotypes were identified using omics approaches (i.e., metabolomics, redox-proteomics, proteomics) and cellular characteristics were analyzed using multiple assays (i.e., deformability using a spleen-mimetic filter, adhesion properties, phosphatidylserine exposure, osmotic resistance, intracellular ATP concentration, proteasome activity). Finally, their circulatory capabilities were evaluated using an *ex vivo* human spleen perfusion model.

## Results

### CFSE^high^ RBCs are storage-induced micro-erythrocytes that can be sorted by flow cytometry

CFSE^high^ RBCs were quantified weekly throughout storage in 8 RBC concentrates collected from healthy human donors and stored in SAGM solution. For all donors, mean CFSE^high^ RBC (min-max ± SEM) accumulation during storage ranged from 1.0% (0.3%-2.8% ± 0.3 %) on day 7 to 31.1% (11.6%-45.2% ± 3.5 %) on day 42, with marked inter-donor variability (Figure 1A). The kinetics and amplitude of CFSE^high^ RBCs accumulation were similar to the SME subset previously quantified by imaging flow cytometry. Quantifying SMEs in the sorted CFSE-stained long-stored RBC subsets (35-42 days of storage) showed that the CFSE^low^ subset contained 4.9% SMEs, whereas the CFSE^high^ subset contained 93.7% SMEs (Figure 1B). Scanning electron microscopy confirmed that sorted CFSE^low^ and CFSE^high^ subsets were highly-enriched in morphologically-normal RBCs (discocytes, echinocytes I and II) and SMEs (echinocytes III, sphero-echinocytes, spherocytes), respectively (Figure 1C). These data confirm that highly-enriched preparations of prone-to-be-cleared SMEs and also of long-stored morphologically-normal *in vitro*-aged RBCs can be obtained, allowing comparison of their properties.

**Figure 1:**
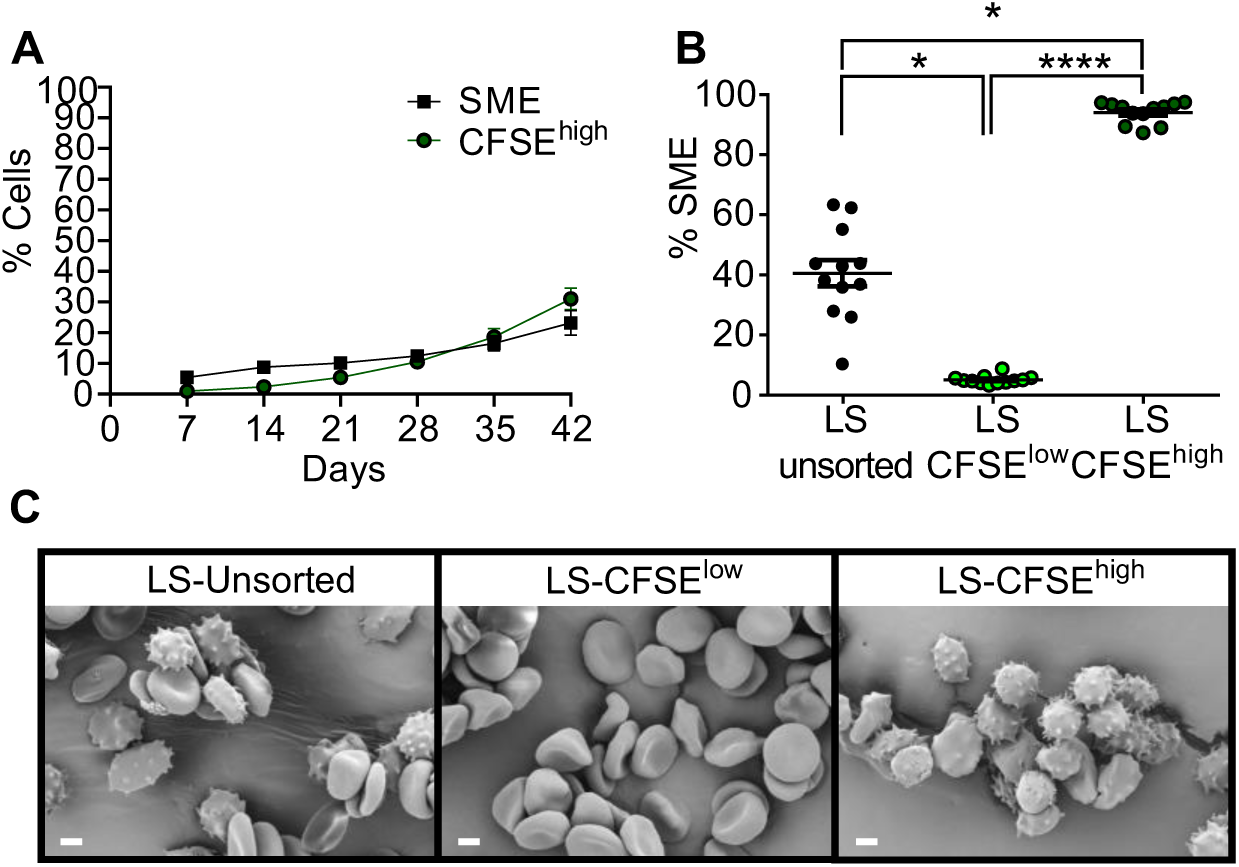
CFSE^high^ RBCs are storage-induced micro-erythrocytes that can be sorted by flow cytometry. **(A)** Weekly quantification of Storage-induced Micro-Erythrocytes (SMEs, black squares) and CFSE^high^ RBCs (green circles) in RBC concentrates stored in SAGM solution at 4°C for 42 days (mean ± SEM of 8 RBC concentrates). **(B)** Proportion of SMEs in CFSE-stained long-stored unsorted (LS-unsorted), and flow-sorted CFSE^low^ (LS-CFSE^low^) and CFSE^high^ (LS-CFSE^high^), RBC subsets. Data are represented as individual points with mean ± SEM of 12 RBC concentrates **(C)** Representative scanning electron microscopy images showing typical RBC morphology of CFSE-stained RBCs. Scale bar represents 2 μm. In panel B, * P < 0.05, **** P < 0.0001 by a Friedman one-way ANOVA followed by Dunn’s multiple comparison test (n = 12).

### Metabolomics identifies subset-specific metabolic alterations in long-stored CFSE^high^ RBCs during aging *in vitro*

Metabolomics analysis compared short-stored CFSE^low^ (stored for 3-12 days) and long-stored CFSE^low^ and CFSE^high^ subsets (stored for 35-42 days). Principal Component Analysis (PCA) illustrates the clear separation of the CFSE^high^ subset from the two CFSE^low^ subsets (Figure 2A). Heatmap representation of the 30 identified metabolites differing significantly between the three subsets also showed this separation, because individual CFSE^high^ samples clustered together, whereas short-stored and long-stored CFSE^low^ RBCs were much more similar to each other (Figure 2B). Indeed, when comparing short-stored and long-stored CFSE^low^ subsets, only 8 metabolites differed significantly, including small variations in energy and redox metabolites for long-stored RBCs (Supplementary Figure 1).

**Figure 2:**
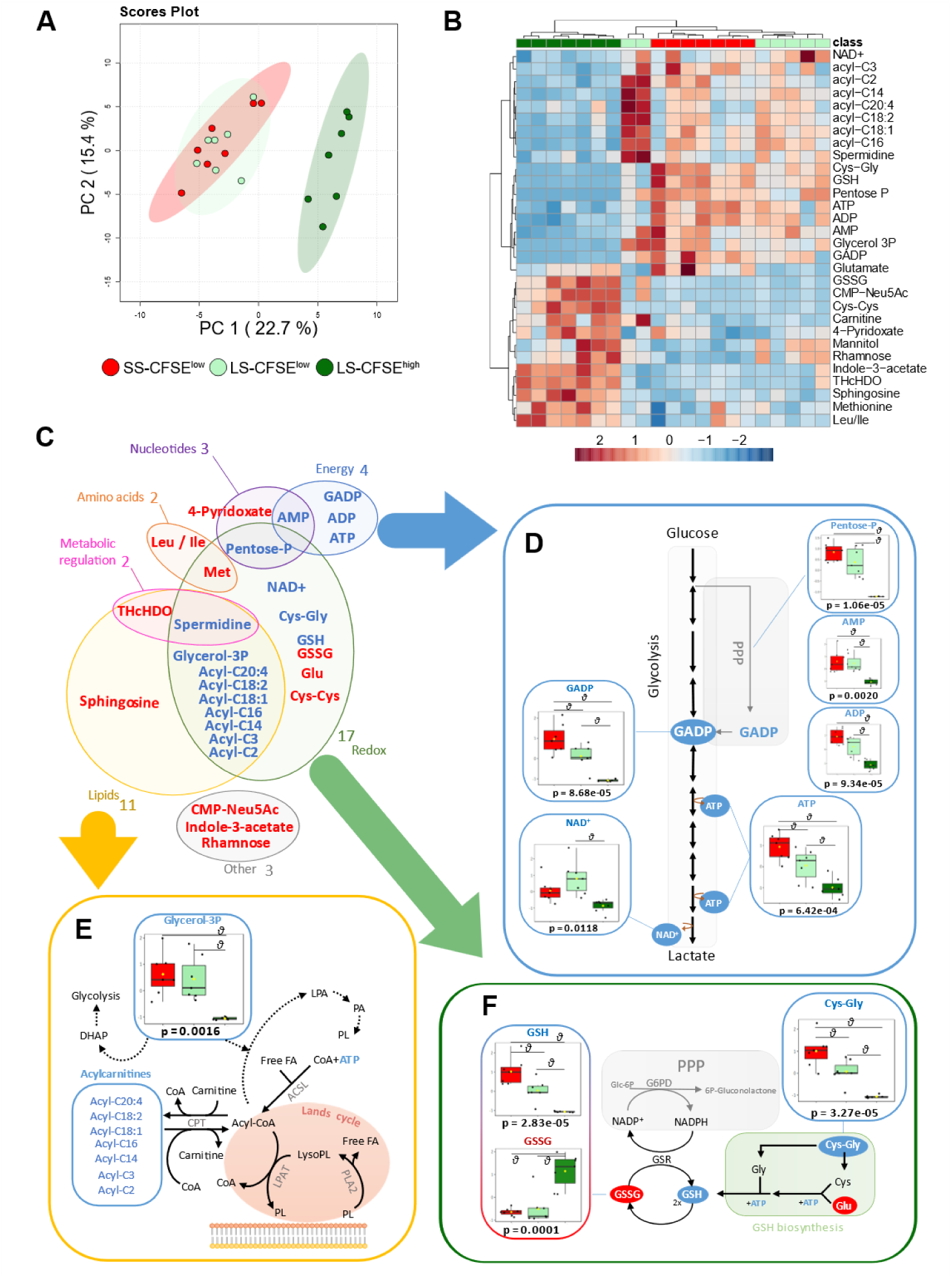
Metabolomics identifies subset-specific alterations in the metabolism of long-stored CFSE^high^ RBCs during aging *in vitro*. **(A)** Principal component analysis of metabolomics data on flow-sorted short-stored CFSE^low^ (SS-CFSE^low^, in red), long-stored CFSE^low^ (LS-CFSE^low^, in light green) and long-stored CFSE^high^ (LS-CFSE^high^, in dark green) RBCs. **(B)** Hierarchical clustering analysis of the 30 metabolites whose levels vary among the three groups by ANOVA followed by Tukey’s multiple comparison test. **(C)** Schematic distribution of the 28 metabolites whose levels vary significantly when comparing long-stored CFSE^low^ and CFSE^high^ RBC subsets. Overview of glycolysis **(D)**, lipid repair **(E)**, and glutathione **(F)** pathways, highlighting key metabolites whose levels vary among the 3 groups. In panels C-F, the metabolites that show significant increases (red font) and decreases (blue font) in long-stored CFSE^high^ RBCs (*vs* long-stored CFSE^low^ RBCs) are shown. Arrows represent a single metabolic step in the pathways and dotted lines represent multiple steps. P-values of ANOVA followed by a FDR correction are indicated under each graph and *θ* represents a significant difference found by a positive post-hoc test of Tukey’s between groups. Acronym definitions are detailed in supplementary data.

However, when comparing long-stored CFSE^low^ and CFSE^high^ subsets, 28 metabolites differed significantly, with major differences in redox, lipid, nucleotide, amino acid, energy, and metabolic regulation (Figure 2C). For energy metabolism (Figure 2D), significant decreases in steady-state metabolite levels in the glycolytic and pentose phosphate pathways were observed in long-stored CFSE^high^ RBCs, including hexose-phosphate (isomers), glyceraldehyde-3-phosphate (GADP), NAD+, and pentose phosphate (isomers), accompanied by decreases in the total adenylate pool, comprising high energy adenosine tri-, di-, and mono-phosphate (ATP, ADP, AMP). Regarding lipid metabolism, glycerol-3-phosphate (Glycerol-3P; implicated in energy and lipid processes) was significantly decreased, along with 7 different acyl-carnitines that participate in lipid recycling through the Lands cycle (Figure 2E, Supplementary Figure 2). For redox metabolism, oxidized metabolites significantly increased, including glutathione disulfide (GSSG), cysteinylcysteine (Cys-Cys; an oxidized form of cysteine), and glutamate (Glu)), accompanied by very low levels of NAD+, reduced glutathione (GSH), and its precursor, cysteinylglycine (Cys-Gly; Figures 2C and 2F). These data suggest that the metabolic storage lesion that occurs during aging *in vitro*, which was previously identified at the whole-population level, is distributed unevenly among individual RBCs and is concentrated in the population of SMEs.

### Redox proteomics identifies a subset-specific capability to resist storage-induced oxidative stress during aging *in vitro*

Post-translational redox modifications were quantified for reversible oxidation (especially of cysteine and methionine) and irreversible oxidation (e.g., cysteine converted to dehydroalanine via beta-elimination of thiols) in short-stored CFSE^low^ and long-stored CFSE^low^ and CFSE^high^ subsets. Unsupervised PCA illustrates the impact of storage on reversible oxidations in both long-stored CFSE^low^ and CFSE^high^ subsets, as compared to short-stored CFSE^low^ RBCs (Figure 3A, left panel). However, no global differences were observed for irreversible oxidations (Figure 3A, right panel). Significant reversible oxidations were detected for 77 proteins, whereas 6 proteins were irreversibly oxidized.

**Figure 3:**
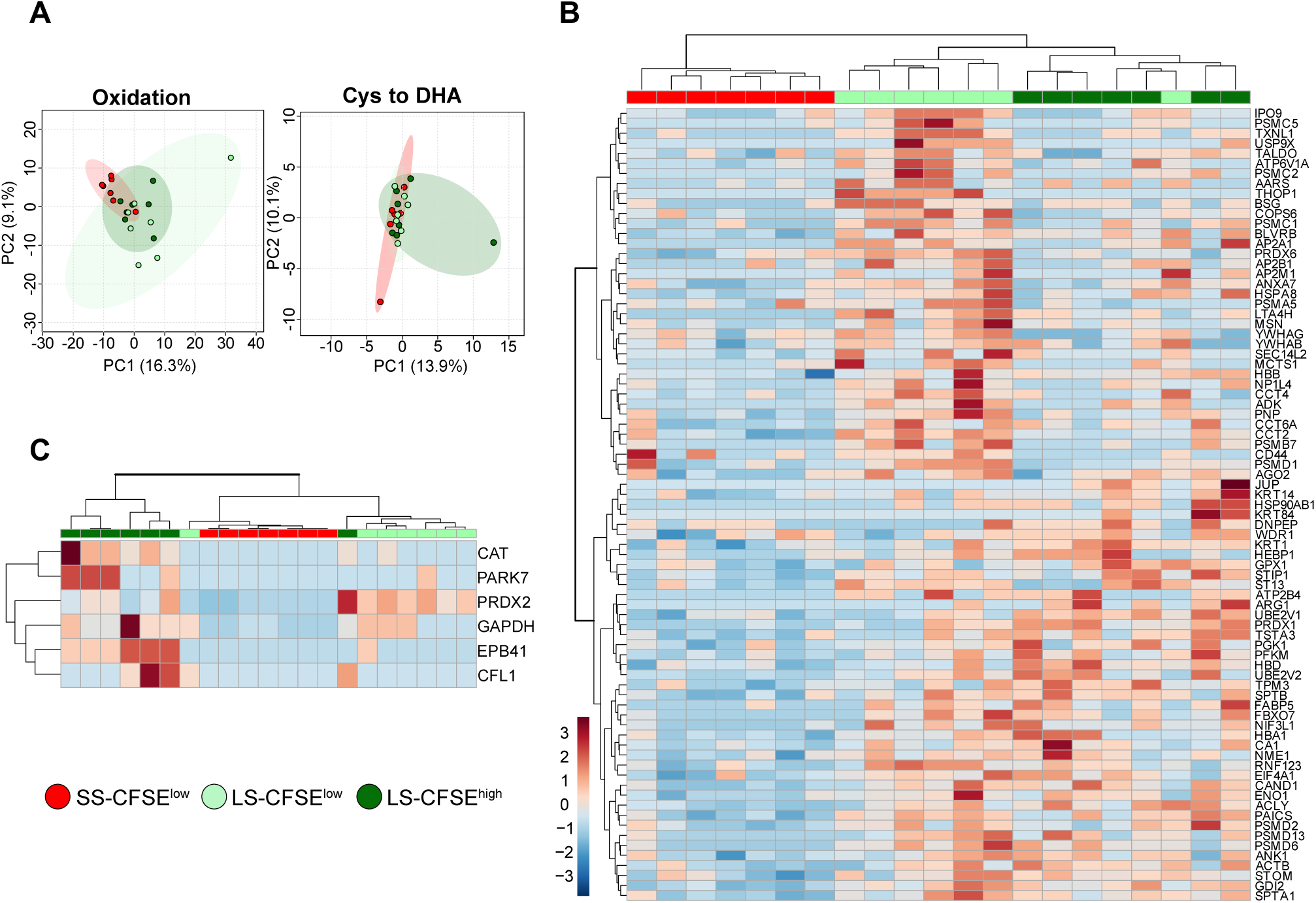
Redox proteomics identifies a subset-specific capacity to resist storage-induced oxidative stress during aging *in vitro*. **(A)** Principal component analysis of reversible (“oxidation”) and irreversible (“Cys to DHA”) protein oxidation data obtained from flow-sorted short-stored CFSE^low^ (SS-CFSE^low^, in red), long-stored CFSE^low^ (LS-CFSE^low^, light green) and long-stored CFSE^high^ (LS-CFSE^high^, in dark green) RBCs. **(B)** Hierarchical clustering analysis of the 79 proteins (out of a total of 659 proteins analyzed, representing 11.7%) with significant reversible oxidation across the three groups by ANOVA followed by Tukey’s multiple comparison test. **(C)** Hierarchical clustering analysis of the 6 proteins (out of a total of 176 proteins analyzed, representing 3.6%) with significant irreversible oxidation across the three groups by ANOVA followed by Tukey’s multiple comparison test.

Heatmap representation of the 77 proteins with reversible oxidations confirmed the impact of storage, but also revealed subset-specific oxidized proteins when comparing long-stored CFSE^low^ and CFSE^high^ RBCs (Figure 3B). Reversibly oxidized proteins (Supplementary Table 1) derive from 6 main families: proteostasis (29%), cytoskeleton (11%), hemoglobin (3.6%), antioxidant system (8%), transport (6%), and glycolysis (6%).

By heatmap representation of the 6 irreversibly oxidized proteins, most CFSE^high^ samples clustered together, whereas short-stored and long-stored CFSE^low^ subsets were much more similar (Figure 3C). The irreversibly oxidized proteins (Supplementary Table 1) involve the cytoskeleton (CFL1 and EPB41), glycolysis (GAPDH), and antioxidant enzymes (CAT, PARK7, PRDX2). Among these, increased irreversible oxidation of four proteins was only detected in the CFSE^high^ subset (CAT, PARK7, CFL1, EBP41). Two proteins were irreversibly oxidized in both long-stored CFSE^high^ and CFSE^low^ subsets (PRDX2, GAPDH), consistent with previous observations (17, 42, 43). Thus, irreversible protein oxidation mainly affected CFSE^high^ RBCs, suggesting that SMEs have a reduced capability to handle and repair storage-induced oxidative stress.

### Proteomics identifies a subset-specific membrane relocation of proteins in long-stored CFSE^high^ RBCs, notably impacting proteins of the proteostasis family

Proteomics analysis compared short-stored CFSE^low^ RBCs with long-stored CFSE^low^ and CFSE^high^ subsets. Proteomics of whole, intact RBCs revealed no significant differences between these three groups (Supplementary Figure 3). However, proteomics of the isolated RBC plasma membranes (i.e., “ghosts”) identified major differences between these three groups. By PCA, the long-stored CFSE^high^ subset is clearly separated from the CFSE^low^ subsets (Figure 4A). Heatmap clustering confirmed the uniqueness of CFSE^high^ RBCs (Figure 4B), revealing expression differences of 96 proteins between the long-stored RBC subsets. However, no significant differences were seen when comparing membrane proteomes of short- and long-stored CFSE^low^ subsets.

**Figure 4:**
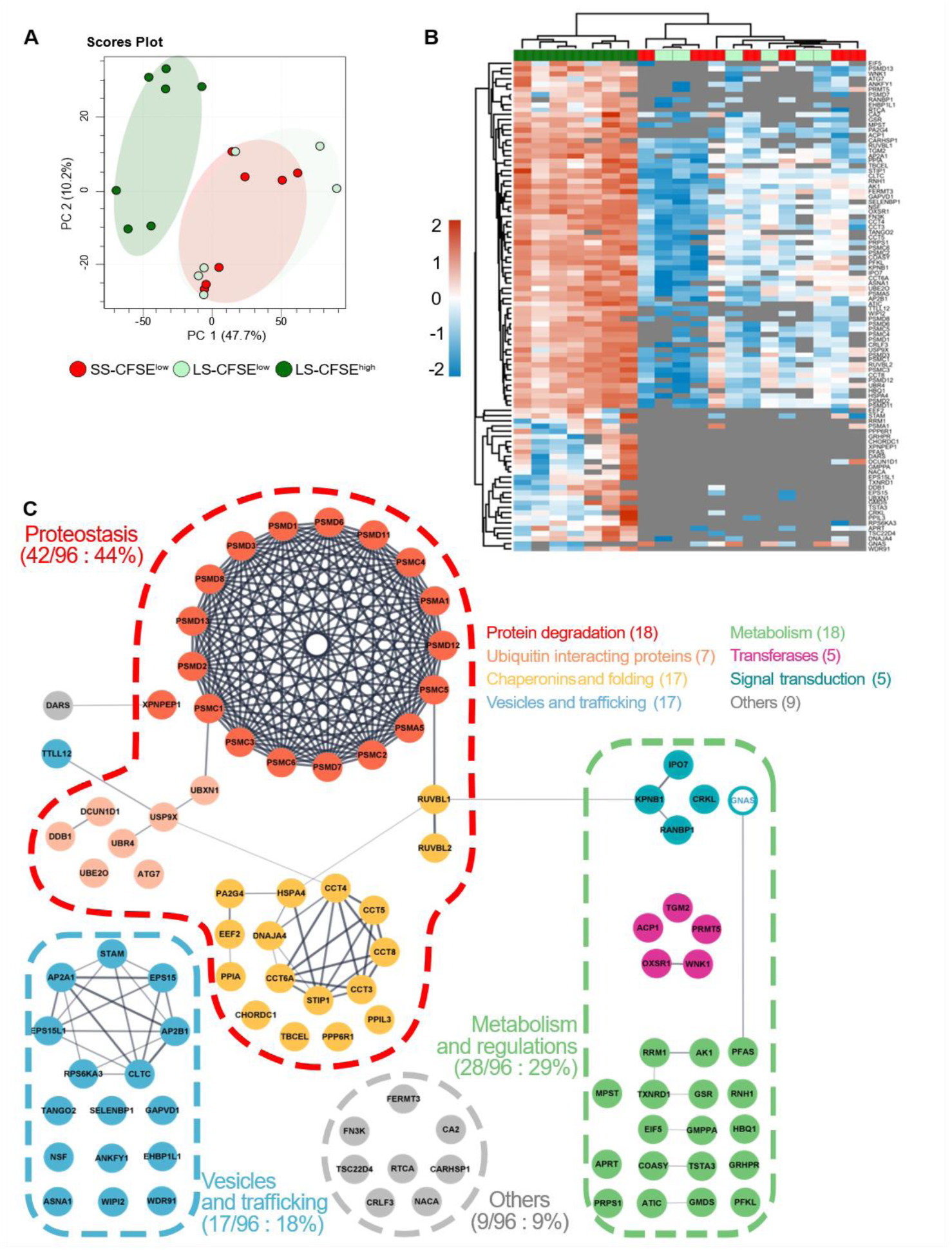
Proteomics identifies subset-specific membrane relocation of specific proteins in long-stored CFSE^high^ RBCs, notably affecting proteins in the proteostasis family. **(A)** Principal component analysis of proteomics data obtained from membrane preparations isolated from flow-sorted short-stored CFSE^low^ (SS-CFSE^low^, in red), long-stored CFSE^low^ (LS-CFSE^low^, light green) and long-stored CFSE^high^ (LS-CFSE^hig^, dark green) RBCs. **(B)** Hierarchical clustering analysis of the 96 proteins that significantly differ between the 3 groups following Pearson clusterization and a FDR correction without data imputation (proteins detected in at least 70% of samples were considered). A z-score scale calculated from copy number/cell shows each protein level while lack of detection is represented by a grey area. **(C)** Interaction network analysis for proteins increased (full filled circle) and decreased (white, transparent, inner circle) in membrane preparations of long-stored CFSE^high^ RBCs, as compared to long-stored CFSE^low^ RBCs. Grey lines represent physical interactions between proteins. This network was realized using Cytoscape StringApp 3.9. Colored dotted lines represent the main protein families identified (relative proportion within the 96 significant proteins) and colored circles represent functional groups within a family.

A physical interaction network (Figure 4C) illustrates potential interactions between the overrepresented proteins in CFSE^high^ RBCs (95/96 proteins). Three protein families were predominantly affected: Proteostasis (44%), Metabolism and regulations (29%), and Vesiculation and trafficking (18%). The increased levels of 95 proteins in long-stored CFSE^high^ membranes, along with their stable proteome at the intact RBC level, suggests that these proteins were relocated to the plasma membrane in this subset.

The most represented family (i.e., Proteostasis) comprises 18 proteins involved in protein degradation, including most 19S proteasome subunit proteins, 17 proteins involved in protein folding, and 7 ubiquitin interacting proteins. Quantitative analysis of copy numbers from intact RBC and membrane proteomics data for proteostasis proteins revealed massive relocation to the membrane of long-stored CFSE^high^ RBCs of most 19S proteasome proteins (mean±SEM; 77±29%; Supplementary Figure 4A), whereas 20S and 11S proteasome proteins showed minimal relocation (5±3% and 9±1%, respectively; Supplementary Figures 4B and 4C, respectively). Ubiquitin interacting proteins showed variable membrane relocation, reaching >50% for 3 proteins (ATG7, UBN4, UBXN1; Supplementary Figure 4D). Most HSP60 and HSP40 chaperone proteins also showed significant membrane relocation (30±8% and 34±27%; Supplementary Figures 5A and 5B, respectively), whereas selected proteins among the HSP70, HSP90, and other chaperone families exhibited slight membrane relocation in long-stored CFSE^high^ RBCs (Supplementary Figure 5C-E). Taken together, these observations implicate the inability of SMEs to maintain proper energy and redox metabolism during aging *in vitro* and also suggest that proteostasis is impaired in these cells.

### Proteasomal degradation capability decreases during RBC storage, especially in CFSE^high^ RBCs

Proteostasis is a protein quality control process that safeguards the cellular proteome by stabilizing correctly-folded proteins, refolding misfolded proteins, and degrading oxidized/misfolded proteins (44). Proteasomal degradation of oxidized/misfolded proteins contributes to cell proteostasis; interestingly, in previous studies, RBC cytosolic proteasome activities gradually decreased during storage (45, 46). We confirmed a slight non-significant decrease in chymotrypsin-like (16%), trypsin-like (7%), and caspase-like activities (19%) of the proteasome during storage (Figure 5A). More importantly, by comparing long-stored CFSE^high^ and CFSE^low^ RBCs, significant decreases in chymotrypsin-like (92%, p<0.01), trypsin-like (75%, p<0.01), and caspase-like (73%, p<0.01) proteasome activities were identified in the CFSE^high^ subset (Figure 5B-5D). In contrast, long-stored and short-stored CFSE^low^ RBCs had similar proteasome activity levels (Supplementary Figure 6). Taken together, these results identify impaired proteasome degradation capability in SMEs, confirming altered proteostasis function in this morphologically-altered subset that increases during aging *in vitro*.

**Figure 5:**
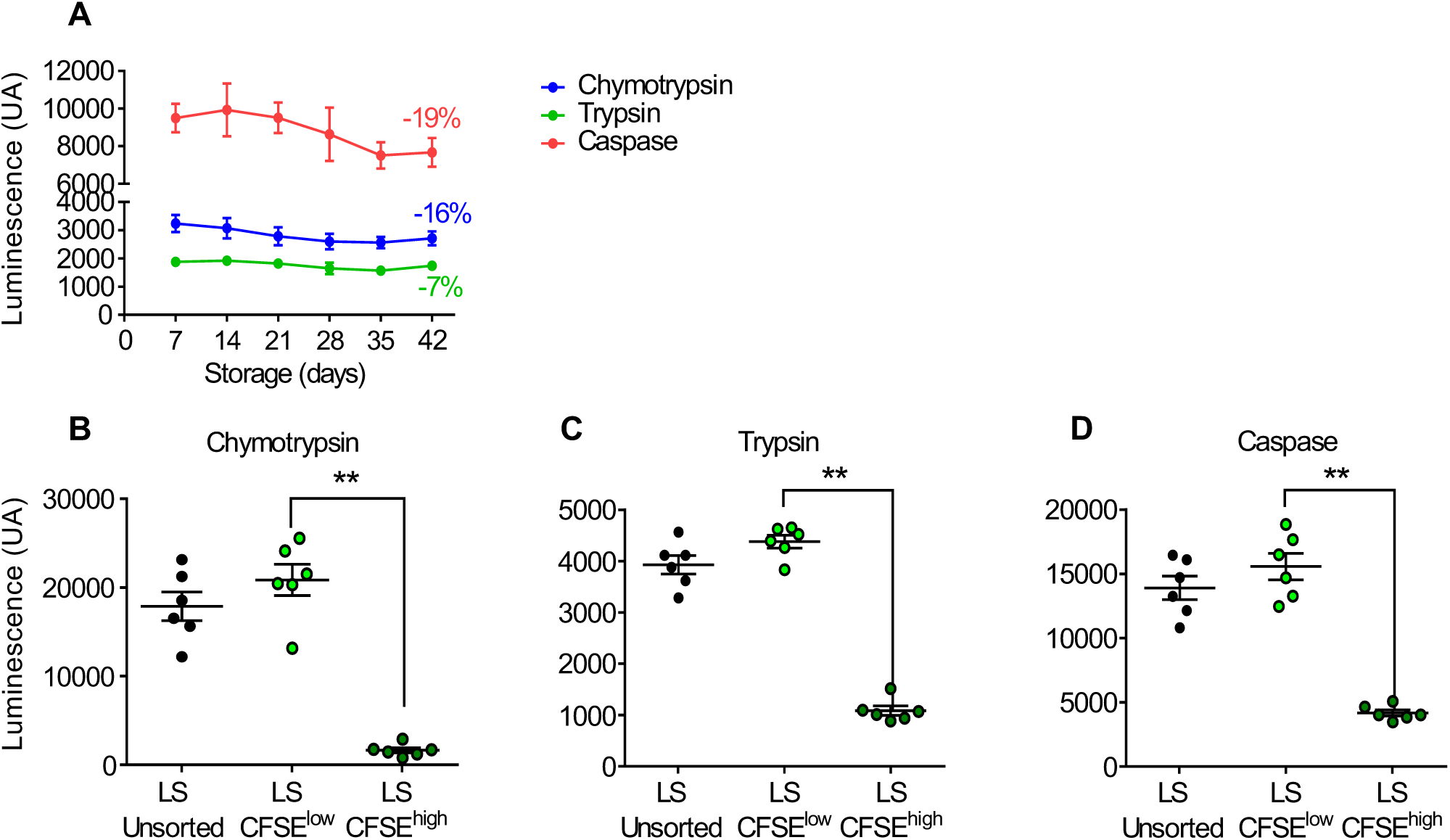
Proteasomal degradation capacity decreases during storage, especially in CFSE^high^ RBCs. **(A)** Weekly quantification of the chymotrypsin-like (blue curve), trypsin-like (green curve), and caspase-like (red curve) proteasome activities during storage of RBC concentrates in SAGM solution for 42 days (mean ± SEM of 7 RBC concentrates). Chymotrypsin-like **(B)**, trypsin-like **(C)** and caspase-like **(D)** activities were measured on subsets for CFSE-stained long-stored unsorted (LS-unsorted), and flow-sorted CFSE^low^ (LS-CFSE^low^) and CFSE^high^ (LS-CFSE^high^), RBCs. In Panel A, a two-way ANOVA followed by Dunnett’s multiple comparison test was performed and no significant difference was observed. In Panels B-D, data are represented as individual points with mean ± SEM of 6 RBC concentrates and ** P <0.01 by Friedman one-way ANOVA followed by Dunn’s multiple comparison test.

### Storage lesions occurring during aging *in vitro* are concentrated in long-stored CFSE^high^ RBCs, which are preferentially cleared during *ex vivo* human spleen perfusion

Storage lesion cell biological characteristics were compared between long-stored CFSE^high^ and CFSE^low^ subsets. Their deformability was evaluated by microsphiltration, an *in vitro* spleen-mimicking device (47). The mean retention rate of CFSE^high^ RBCs was significantly increased (27.5%), as compared to CFSE^low^ RBCs (-10.6%), indicating reduced deformability of CFSE^high^ RBCs (p<0.05, Figure 6A). Dynamic endothelial cell adhesion experiments showed a significant 11-fold increase in adherence of long-stored CFSE^high^ RBCs (1720 RBC/cm²), as compared to CFSE^low^ RBCs (154 RBC/cm², p<0.001, Figure 6B). PS-exposing RBCs were identified by lactadherin binding. Since lactadherin-FITC detection by flow cytometry is not compatible with CFSE staining, CTV staining was used to sort CFSE^high^ and CFSE^low^ subsets using a different flow cytometry channel (Supplementary Figure 7). The proportion of PS-exposing long-stored RBCs in the CTV^high^ subset (14.5%) was increased, as compared to the CTV^low^ subset (0.3%, p<0.01, Figure 6C). Resistance to osmotic hemolysis also decreased in long-stored CFSE^high^ RBCs (0.52%), as compared to CFSE^low^ RBCs (0.47%, p<0.01, Figure 6D). Finally, intracellular ATP levels in long-stored CFSE^high^ RBCs were very low (0.12 µmol/g Hb), and significantly less (p<0.0001) than in CFSE^low^ RBCs (4.69 µmol/g Hb, Figure 6E). Interestingly, long-stored and short-stored CFSE^low^ RBCs had similar intracellular ATP levels (4.69 *vs* 5.58 µmol/g Hb, respectively; Supplementary Figure 8). Taken together, these results show that cellular storage lesions are concentrated in the SME subset during aging *in vitro*.

**Figure 6:**
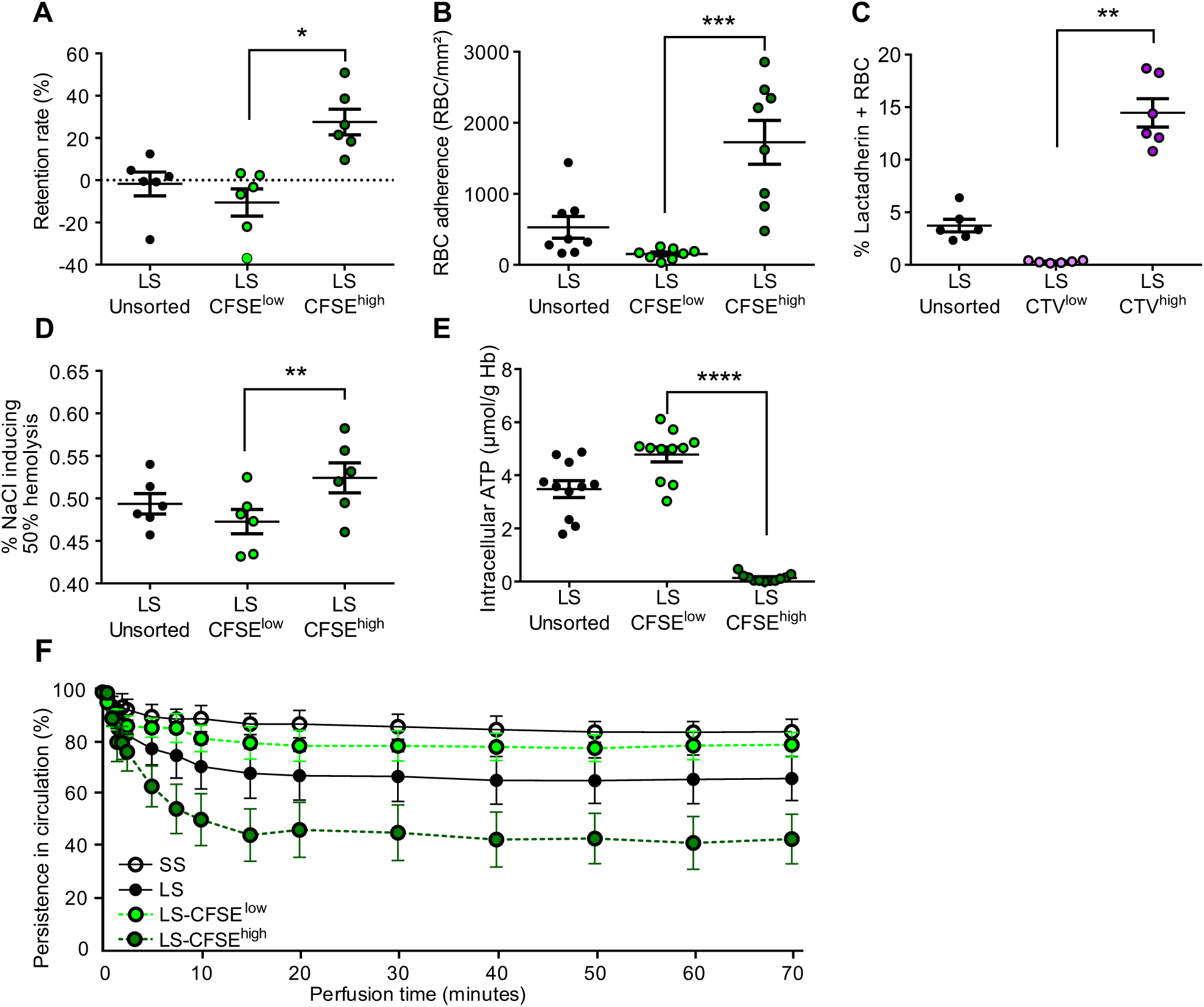
Storage lesions occurring during aging *in vitro* are concentrated in long-stored CFSE^high^ RBCs, which are preferentially cleared during *ex vivo* human spleen perfusion. **(A)** Retention rate by microsphiltration, **(B)** dynamic RBC adhesion to endothelial cells, **(C)** phosphatidylserine (PS) exposure quantified by lactadherin staining, **(D)** osmotic fragility determined by measuring the NaCl concentration required to induce 50% hemolysis, **(E)** intracellular ATP levels normalized to hemoglobin content were measured on CFSE-stained unsorted long-stored RBCs (LS-unsorted) RBCs, and on flow-sorted LS-CFSE^low^ and LS-CFSE^high^ RBCs. **(F)** Kinetics (mean ± SEM of 4 independent perfusions) of the normalized circulating concentrations of stained short-stored (SS), long-stored (LS), LS-CFSE^low^, and LS-CFSE^high^ RBC subsets during *ex vivo* perfusion of a human spleen. In **(C)**, CTV-stained RBCs were analyzed to allow for staining using lactadherin-FITC. In Panels A-E, data are represented as individual points with mean ± SEM of ≥6 RBC concentrates). * P <0.05, ** P <0.01, *** P <0.001, **** P <0.0001 by Friedman one-way ANOVA followed by Dunn’s multiple comparison test.

Finally, we compared the circulatory capability of long-stored RBCs mixed with short-stored RBCs using *ex vivo* perfusion of human spleens (4 independent experiments, Figure 6F). The mean proportion of circulating short-stored and long-stored RBCs decreased by 15% and 33%, respectively, over 70 minutes, confirming that long-stored RBCs have decreased circulation capability as compared to short-stored RBCs. Among long-stored RBCs, the two CFSE subsets had very distinct profiles of circulatory persistence, with the mean proportion of CFSE^high^ RBCs decreasing by 57%, whereas that of CFSE^low^ RBCs decreased by only 20%; the latter was similar to that found with short-stored RBCs. These results confirm that short-stored and long-stored morphologically-normal CFSE^low^ RBCs are similar in their capability to circulate in this *ex vivo* perfusion model of transfusion. They also suggest that the molecular and cellular alterations of SMEs that were identified herein induce their clearance in this *ex vivo* transfusion model.

## Discussion

The components of the RBC storage lesion occurring during aging *in vitro* are typically assessed at the whole-population level; as such, any particular result is averaged out over the entire population of RBCs in a given volunteer donor unit. By employing a staining protocol to specifically sort long-stored morphologically-altered RBCs from morphologically normal RBCs, we show that morphologically-altered CFSE^high^ RBCs are mainly SMEs, which progressively accumulate during storage and are preferentially retained in the spleen. These findings support previous observations characterizing post-transfusion clearance of SMEs in a mouse model and their negative correlation with post-transfusion recovery in healthy human volunteers (40). Molecularly, significant changes in energy, redox, and lipid-repair metabolism mainly occurred in these morphologically-altered RBCs. Additionally, these morphologically-altered RBCs exhibited distinct profiles of proteins that were reversibly oxidized, irreversibly oxidized, and relocated to the membrane (including chaperones and most proteins of the 19S proteasome subunit), accompanied by decreased proteasomal enzymatic activity. At the cellular level, characteristics typically associated with RBC clearance were found with the morphologically-altered RBCs. Conversely, cellular and molecular properties, and lack of splenic retention, of long-stored morphologically-normal RBCs resembled those of short-stored RBCs. Taken together, these findings strongly support the concept that the storage lesion differentially affects individual RBCs, revealing biological networks that are overwhelmed by metabolic and oxidative stresses in a specific RBC subset particularly sensitive to the *in vitro* aging process.

Our enzymatic assay and metabolomics data confirm the well-documented decrease in intracellular ATP levels during RBC storage (6–8, 48, 49), and further document that the morphologically-altered RBCs are more severely affected than morphologically-normal RBCs. ATP is required by numerous enzymes, including flippases that internalize PS, and membrane pumps that maintain cellular homeostasis; thus, ATP is essential for RBC survival (50–53). In addition, ATP levels negatively correlate with post-transfusion recovery (7, 36, 37, 39, 54), and its subset-specific depletion in SMEs could trigger a cascade of events inducing senescence and clearance.

Depleted pentose phosphates and GAPD metabolites in SMEs identify subset-specific dysregulation of the pentose phosphate pathway (already observed at the whole-population level) (1, 17, 55), suggesting inadequate NADPH levels that are necessary to support multiple antioxidant pathways (56–58), including recycling oxidized glutathione to its reduced form by glutathione reductase. Interestingly, we identified subset-specific depletion of reduced glutathione (GSH) and its precursor (cysteinylglycine), and massive increases in oxidized glutathione (GSSG), suggesting significant GSH consumption by redox-sensitive thiols (59, 60). Because glutathione synthesis is also ATP-dependent, depleting this important antioxidant system (61, 62) hampers the resistance of SMEs to storage-induced oxidative stress.

Depletion of acylcarnitine metabolites also suggests that the mechanisms required to repair oxidatively-damaged membrane lipids are impaired in SMEs. Indeed, lipid repair through the Lands cycle is fueled by the acylcarnitine pool in RBCs (63). Interestingly, genetic control of carnitine metabolism was recently shown to determine hemolytic propensity during aging *in vitro* and *in vivo* (64), and impaired lipid repair mechanisms are also observed in sickle cell disease (65).

As RBCs age during storage, and protective systems fail, oxidant stress induces protein oxidations that are initially reversible, and then become irreversible, such as with carbonylation (5, 12, 13, 15, 16) and with beta-elimination of cysteine thiol groups to generate dehydroalanine (17, 60). Our data confirm storage-dependent reversible oxidation in both SMEs and morphologically-normal long-stored RBCs, as compared to short-stored RBCs. However, irreversible beta-elimination of thiol groups was mainly detected in SMEs, supporting the finding of decreased glutathione antioxidant protection in this subset that is particularly sensitive to the *in vitro* aging process.

Proteomics revealed massive relocation of cytosolic proteins to the membrane of morphologically-altered RBCs. Oxidized/misfolded proteins tend to form cytotoxic hydrophobic aggregates, removing them from the pool of active molecules, thereby reducing their function (66). Due to their ability to recognize and bind misfolded/oxidized proteins, proteostasis components are particularly prone to becoming sequestered in aggregates. In this case, they are substantially depleted from the soluble, active pool, negatively impacting the proteostasis network by limiting its capacity, thereby contributing to a vicious circle leading to further aggregate accumulation (67–71). This is particularly relevant for mature RBCs, which no longer have the ability to synthesize new proteins *de novo*. Our observation that proteostasis proteins relocate to the RBC membrane strongly supports the concept that proteostasis functions are hampered in the morphologically-altered RBCs.

Oxidized proteins are susceptible to ATP-independent proteasomal degradation or ubiquitination by ubiquitin-interacting proteins, which then targets them for ATP-dependent proteasomal degradation (72, 73). Our results show that proteasomal enzymatic activity is decreased in SMEs, suggesting that the mild decrease in proteasome activity observed during storage (at the whole-population level) (45, 46) is due to significant, subset-specific decreases in SMEs. Decreased proteasome activity could be due to ATP depletion, proteasome component relocation to the membrane and/or release in vesicles (45, 46), and/or inefficient unfolding of oxidized/misfolded proteins, preventing their access to the proteasome catalytic core (15, 74).

The relocation of proteins and protein aggregates to the RBC membrane induces irreversible shedding of RBC cytosolic and membrane constituents by vesiculation, resulting in the formation of SMEs through membrane loss, and leading to the accumulation of microparticles in storage bag supernatant (75–78). Thus, loss of proteostasis function, accompanied by vesiculation, are consistent with the RBC subset-specific morphological alterations that produce SMEs. The cellular alterations of the morphologically-altered RBCs observed herein, including increased PS exposure (a pro-phagocytic “eat me” signal) (79), endothelial cell adhesion, retention in a spleen-mimetic filter (consistent with diminished deformability from a lower surface/volume ratio) (80, 81), and osmotic fragility (82), could all contribute to RBC retention in the *ex vivo* spleen perfusion model (40, 83).

Our data show that only a subset of RBCs is severely altered during storage; the available literature suggests that this corresponds to the older RBCs collected at donation (84, 85). Therefore, our data support the concept that the RBC storage lesion is a form of aging *in vitro* that enhances aging processes already underway in *in vivo*. For example, during aging *in vivo*, low ATP levels are seen in senescent RBCs and, because redox capacity depends upon energy metabolism, they may be more vulnerable to oxidative stresses encountered during refrigerated storage (86, 87). In addition, proteostasis dysfunction is a known hallmark of aging nucleated cells; we now propose that it is also important in aging of nucleus- and mitochondria-free RBCs (44). Finally, in response to protein oxidation, the only alternatives that mature, organelle-free RBCs have are to protect, repair, destroy, or sacrifice their misfolded/oxidized proteins, since they can no longer synthesize new proteins (88).

In this conceptual framework, our metabolomics, redox proteomics, proteomics, proteasome activity, and cell biological data are all consistent with a progressive loss of proteostasis function in the older RBCs present “in the bag” at the time of blood donation, leading to the accumulation of SMEs during storage, which are cleared by the recipient’s spleen post-transfusion, thereby linking proteostasis dysfunction to RBC clearance (Figure 7).

**Figure 7:**
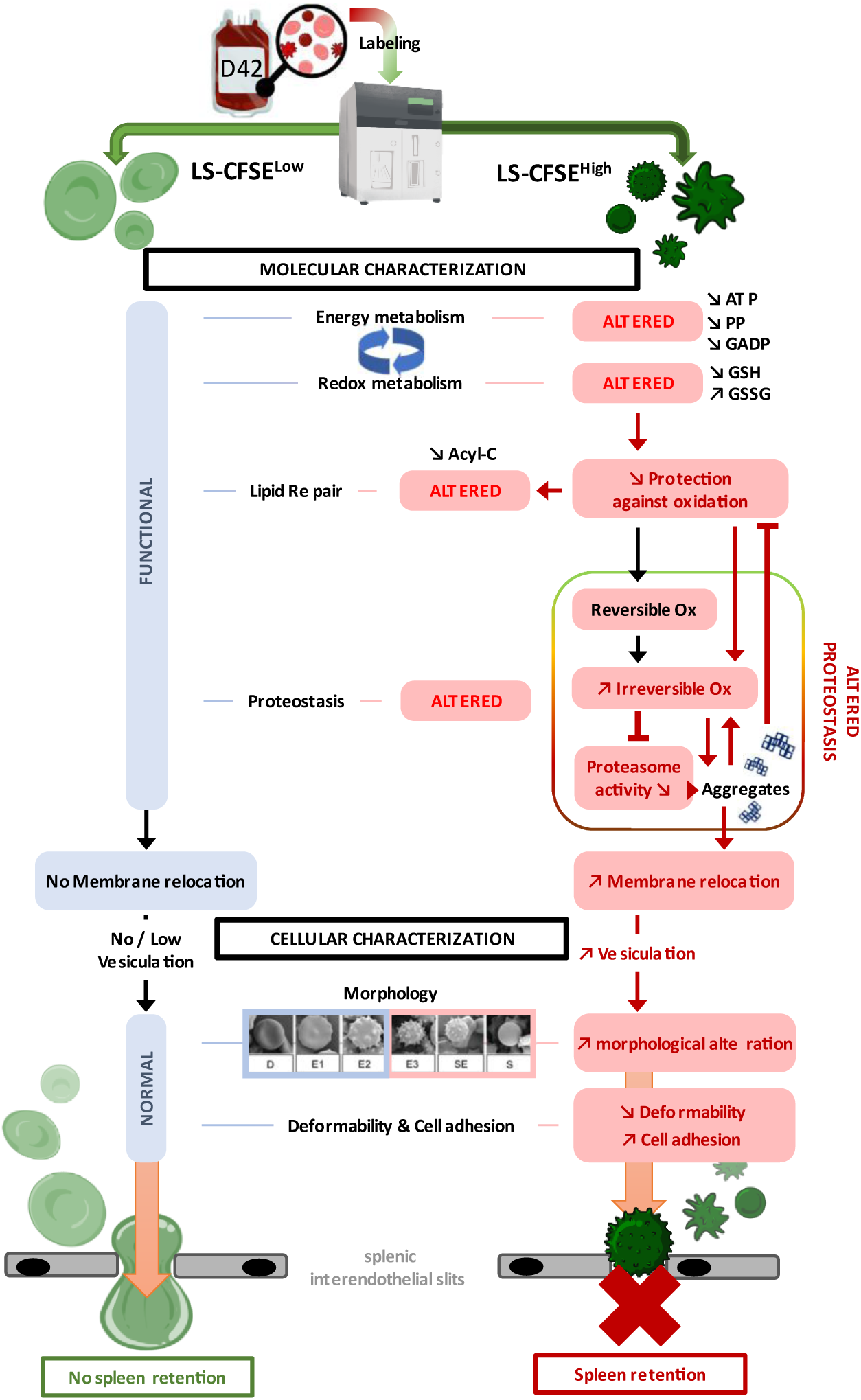
A proposed model to illustrate the main alterations affecting each RBC subset during storage. The main results of the comparative molecular and cellular characterizations (black boxes) of long-stored CFSE^low^ (light green, discocytes) and long-stored CFSE^high^ (dark green, echinocytes III, spheroechinocytes and spherocytes) RBCs. In long-stored CFSE^low^ RBCs (left side), functional energy and redox metabolism sustains effective lipid-repair and proteostasis functions, limiting protein aggregation, membrane relocation, and vesiculation. Functional molecular properties contribute to maintaining normal cellular properties (e.g., morphology, deformability, endothelial cell adhesion) and, thus, their ability to circulate. In long-stored CFSE^high^ RBCs (right side), decreased energy metabolism (e.g., ATP, PP, GAPD) is unable to fuel the redox system effectively, leading to decreased GSH and increased GSSG levels, culminating in dysfunction in lipid repair and proteostasis. The accumulation of irreversibly oxidized proteins could also lead to decreased proteasome function. The combined effect of oxidized protein accumulation and decreased proteasome degradation could fuel a negative feedback loop that further inhibits the redox system, thereby favoring production of toxic protein aggregates, protein relocation to the RBC membrane, and, ultimately, vesiculation. Vesiculation alters morphology, leading to decreased deformability, increased endothelial cell adhesion, and splenic retention. Non-significant (light blue boxes) and significant alterations are shown (pink boxes), black and red lines represent respectively normal and negative interactions. PP: Pentose Phosphate, GADP: DL-Glyceraldehyde-3-phosphate; GSH: reduced glutathione, GSSG: oxidized glutathione, Acyl-C: acyl-carnitines; Reversible – Irreversible Ox: reversible or irreversible oxidation.

Our study has some limitations. For example, the staining protocols require a 6-hour minimum incubation time at 37°C to discriminate between CFSE^high^ and CFSE^low^ subsets. The observed differences may reflect end-of-storage RBC properties, which are normally evaluated directly following storage at 4°C, and/or RBC properties similar to those described in prior *in vitro* transfusion models (89, 90); thus, the observed phenotype may be exacerbated by the 37°C incubation. Despite potential effects of the staining/sorting protocol, the short-stored CFSE^low^ and long-stored CFSE^high^ and CFSE^low^ subsets experienced the same treatment and, therefore, are directly comparable. As another limitation, omics experiments were conducted on a limited number of random donors; further confirmation with additional donors would strengthen our conclusions. Also, regarding omics data, the measured metabolites and proteins represent only a snapshot of cellular activity; performing dynamic flux experiments would confirm and extend the current observations.

The morphologically-altered RBCs that accumulate during storage are preferentially retained in the spleen upon transfusion, and their rapid clearance post-transfusion likely explains the decreased hemoglobin increment seen with long-stored RBCs (25, 34, 35). Quantifying them in RBC concentrates by either imaging flow cytometry (i.e., SMEs) or flow cytometry (i.e., CFSE^high^ RBCs) could provide an *in vitro* marker to predict transfusion efficacy, which could be used to evaluate new manufacturing processes (e.g., hypoxic (39), pathogen-inactivated (91), and DEHP-free storage (92)), storage alternatives (e.g., AS-7 (93)), and donor factors that affect storage quality (25, 34, 94).

Refrigerated storage of organelle-free, hemoglobin-rich RBCs, has historically been regarded as a useful model of oxidant stress biology and cellular aging. The current results confirm these assumptions and suggest that it may also provide a model of proteostasis dysfunction. Additional exploration of proteostasis during RBC storage, and in physiologic and pathophysiologic contexts, could further deepen our understanding of RBC aging *in vitro* and *in vivo*.

## Methods

Detailed information is provided in Supplementary Data.

### Sex as a biological variable

Red blood cells and spleens used in this study are derived from anonymized human samples. Sex was not considered as a biological variable.

### RBC concentrate collection and storage

Leukoreduced RBC concentrates from healthy donors were obtained from the Etablissement Français du Sang and stored in SAGM at 2-6°C for 44 days. Samples were aseptically collected and analyzed on storage day 3-10 (short-stored) and 40-44 (long-stored).

### Carboxyfluorescein diacetate succinimidyl ester (CFSE) and CellTrace Violet (CTV) staining

The staining protocol was performed, as described (41). Briefly, RBCs were stained with 0.05µM CFSE for 20 minutes at 37°C, washed, suspended in RPMI complete (RPMIc; RPMI 1640 with 10% fetal bovine serum (FBS) and antibiotic/antimycotic solution) and incubated overnight at 37°C. CTV staining (1µM) was performed using the same protocol.

### Cell sorting

CFSE^low^ and CFSE^high^ RBCs were sorted using a MA900 Cell Sorter (Sony). Sorted RBCs were centrifuged after collection, resuspended in RPMIc, and stored at 4 °C until analysis or frozen (for omics experiments). As controls, stained RBCs were sorted using only size/structure parameters (unsorted condition).

### Imaging flow cytometry

Imaging flow cytometry (ImageStream X Mark II; Amnis® Flow Cytometry, Luminex) examined RBC dimensions and morphology, as described.^19^ Brightfield images (x60 magnification) were analyzed using dedicated computer software (IDEAS [version 6.2]; Amnis) to determine the proportions of SMEs.

### Scanning Electron Microscopy

RBCs were prepared as described (40) and observed using a Zeiss Ultra plus field emission-scanning electron microscope (ZEISS).

### Metabolomics and redox-proteomics

Metabolomics were performed, as described (95). Redox-proteomics were performed using Filter Aided Sample Preparation digestion and nano ultra-high-pressure liquid chromatography tandem mass spectrometry (17, 96).

### Proteomics of intact RBCs and RBC membranes

Proteomics of intact RBCs and RBC membranes (i.e., ghosts) were performed by nanoscale liquid chromatography coupled to tandem mass spectrometry (nLC-MS/MS), as described (97).

### Phosphatidylserine exposure

Phosphatidylserine (PS)-exposing RBCs were quantified using fluorescein isothiocyanate (FITC)-conjugated bovine lactadherin (lactadherin-FITC, Cryopep), as described (8).

### Microsphiltration

Microsphiltration plates containing calibrated metal microspheres (to mimic spleen inter-endothelial slits) were used to assess RBC deformability, as described (8). Diluent RBCs (95%) were stained with CellTrace Far Red (Life technologies), test RBCs (5%) were sorted CFSE^low^ and CFSE^high^ subsets, and upstream mixtures prepared at a 1% hematocrit in Krebs-albumin solution, filtered through the microsphere layer, and washed twice. Downstream suspensions were then collected. Upstream and downstream proportions of test RBCs were evaluated by flow cytometry and retention rates calculated as: [(UP-DW)/UP] x 100 (UP = % of test RBCs in upstream sample; DW = % of test RBCs in downstream sample).

### ATP

Intracellular ATP concentrations were determined using an ATP assay kit (ATPlite, PerkinElmer) and normalized against the hemoglobin (Hb) concentration of each sample (μmol/g Hb).

### Osmotic fragility

RBC osmotic fragility was determined, as described (19), using 3,3’, 5,5”-tetramethylbenzidine to increase sensitivity.

### Dynamic RBC adhesion to endothelial cells

Dynamic adhesion experiments were performed, as described (8). Briefly, human microvascular endothelial cell line 1 (HMEC-1) cells were cultured in microchannels and RBCs were perfused (10 minutes, 0.2 dyn/cm2). The shear stress was increased every 5 minutes (0.5 and 1 dyn/cm2) to remove less adherent RBCs. Brightfield images at 1 dyn/cm² were analyzed to quantify adherent RBCs.

### Proteasome activity

Proteasome-specific activities (i.e., chymotrypsin-like, trypsin-like, caspase-like) were measured by using Cell-Based Proteasome-Glo™ Assays (Promega), following the manufacturer’s recommendations.

### Human spleen retrieval and ex vivo perfusion

Spleens (macroscopically and microscopically normal) were retrieved and processed, as described (98), from patients undergoing distal splenopancreatectomy for pancreatic disease. The main splenic artery was cannulated and spleens flushed with cold Krebs-albumin solution.

Spleens were perfused with long-stored CFSE-stained RBCs and Celltrace Far Red-stained short-stored RBCs (final hematocrit of 5-30% in Krebs-albumin solution), for 70 minutes at 37°C. Samples were retrieved from the circuit for flow cytometric quantification. Persistence in circulation was calculated as: (% stained RBCs in sample/% stained RBCs at T0) x 100.

### Statistics

Data were analyzed using GraphPad Prism version 9.2.0 for Windows (GraphPad Software). All data with >8 samples were tested for normality with the D’Agostino and Pearson test. ANOVA with Sidak’s multiple comparison test was performed for parametric data. ANOVA of Friedman with Dunn’s multiple comparison test was performed for non-parametric data. A p-value <0.05 was considered statistically significant.

### Study approval

The study was conducted according to the Declaration of Helsinki. Human spleens were retrieved in the context of the Spleenvivo project approved by the “Ile-de-France II” Institutional Review Board on 4 September 2017 (#CPP 2015-02-05 SM2 DC). Written informed consent was received prior to participation.

## Data availability

The values for all data points in graphs are reported in the Supporting Data Values file. Mass spectrometry proteomics raw data were deposited in the ProteomeXchange Consortium via the PRIDE (99) partner repository with dataset identifier PXD049411.

## Author contributions

SP, MM, MD, LL, YH, AF, CR, PN, MKR, MDZ, JB, SG and AS performed experiments. FP and SD provided human spleens. SP, MM, MDU, EFG, ADA and PA analyzed the data. SP, MM and MDU prepared figures. MC, SP, MM, CR, SS, OH, PAB, ADA and PA designed the research. SP, MM, MD and PA wrote the manuscript. CR, EFG, SS, PAB and ADA edited the manuscript. All the authors critically contributed to the progress of the project and finalization of this manuscript.

## Acknowledgments

The authors thank Etablissement Français du Sang « Haut de France-Normandie » and Île-de-France » for providing RBC concentrates. The authors thank Arnaud Chene for his help in setting up the sensitive osmotic fragility test, and Jean-Philippe Semblat and Laetitia Claer for their technical assistance in using the cell sorter. The Orbitrap Fusion mass spectrometer was acquired with funds from the FEDER through the “Operational Program for Competitiveness Factors and employment 2007-2013”, and from the “Cancéropôle Ile-de-France”. SP and MC were supported by a French Ministry of Education and Research and Laboratory of Excellence GR-Ex scholarships, respectively. MD holds a Paris-Cité University research engineer position fully supported by the Laboratory of Excellence GR-Ex. This work was supported by state funding from the Agence Nationale de la Recherche under the “Investissements d’avenir” program (ANR-10-IAHU-01, ANR-11-LABX-0051, and ANR-18-IDEX-0001); and Association Recherche Transfusion (ART) grant 181 and 210. This study was also supported by funds by the National Heart, Lung, and Blood Institute R01HL148151 (SLS, ADA, PA) and R21HL150032, R01HL146442, R01HL149714 (ADA).

