## Supplementary data for "Proteostasis and metabolic dysfunction in a distinct subset of storage-induced senescent erythrocytes targeted for clearance"

### Methods

#### Carboxyfluorescein diacetate succinimidyl ester (CFSE) and CellTrace Violet (CTV) staining

RBCs were washed and stained with CFSE (5.5 million RBC/mL, 0.05µM CFSE in PBS) for 20 minutes at 37°C (1). RBCs were then washed, suspended in RPMIc (RPMI 1640 supplemented with 10% FBS and 1% Antibiotic/Antimycotic solution), and incubated overnight at 37°C. Following incubation, RBCs were washed, resuspended in fresh RPMIc, and stored at 4°C until analysis. CTV staining (1µM) was performed using the same protocol.

#### Cell sorting

Sorting of $CFSE^{low}$ and $CFSE^{high}$ RBCs was performed using a MA900 Cell Sorter (Sony) with a 100µm sorting-chip at the maximum speed of 10,000 events per second in semi-purity mode. Sorted RBCs were collected in tubes containing 1mL of RPMIc, then centrifuged, resuspended in RPMIc, and stored at 4°C until analysis. Stained RBCs that were sorted only based on size/structure parameters were used as controls (unsorted condition).

#### Imaging flow cytometry (IFC) analysis

IFC was performed with an ImageStream X Mark II (Amnis® Flow Cytometry, Luminex, Seattle, WA, USA) to determine RBC morphology (2). RBC were suspended at 1% hematocrit just before acquisition in Krebs-albumin solution (Krebs-Henseleit buffer, Sigma-Aldrich) modified with 2g/L of glucose, 2.1g/L of sodium bicarbonate, 0.175g/L of calcium chloride dehydrate, and 5g/L of lipid-rich bovine serum albumin (Albu-MAX II, Life Technologies). Images (x60 magnification) were recorded (INSPIRE software, AMNIS) using the brightfield channel to be processed using dedicated computer software (IDEAS [version 6.2]; Amnis). Focused cells and single cells were respectively selected using the features gradient

RMS_M01_Ch01 and Aspect ratio_M01_Ch01 versus Area_M01_Ch01. Front views were selected using the feature Circularity_Object (M01, Ch01, Tight) and projected surface area was determined using the feature Area_Object (M01, Ch01, Tight). At least 6000 front views of focused single RBCs/condition were analyzed. The SME proportion was determined independently for each donor, using the nadir of the bimodal frequency histograms as the gating boundary.

**Proteomics of intact RBCs and RBC ghosts**

Proteomics of intact RBCs and RBC ghosts were performed by nanoscale liquid chromatography coupled to tandem mass spectrometry (nLC-MS/MS). The mass spectrometry proteomics data were deposited in the ProteomeXchange Consortium via the PRIDE (3) partner repository with the dataset identifier PXD049411.

<u>Sample processing:</u> To prepare ghosts, hypotonic buffer used to lyse RBCs: 5mM Na2HPO4, 0.35mM EDTA, 1mM phenylmethylsulfonyl fluoride (PMSF). Membranes were pelleted by centrifugation for 20 minutes at 15600g. The ghost pellet was washed several times with hypotonic buffer to obtain a white pellet. Five million RBCs and 10 million ghosts per sample were boiled in lysis buffer (50mM TRIS pH 8.5 with 2% SDS) for 5 min at 95°C. Protein concentrations were determined using a bicinchoninic acid assay (BCA kit; Pierce). Disulfide bridges from 50 micrograms of protein for intact RBC samples, and from the totality of proteins for ghost samples, were reduced using 10mM tris(2-carboxyethyl)phosphine; the resulting free thiols were protected using 50mM chloroacetamide for 5min at 95°C. Proteins were trypsin-digested overnight using the filtered-aided-sample-preparation method, as described (4). Eluted peptides were separated into 5 fractions using strong cation exchange (SCX) StageTips and then vacuum-dried during centrifugation in a Speed Vac (Eppendorf).

nLC-MS/MS proteomics: MS analyses were performed on a Dionex U3000 RSLC nano-LC system coupled to an Orbitrap Fusion mass spectrometer (Thermo Fisher Scientific). Peptides from each SCX fraction were solubilized in 0.1% trifluoracetic acid containing 10% acetonitrile, loaded, concentrated, and washed on a C18 reverse phase precolumn (3-µm particle size, 100 Å pore size, 75-µm inner diameter, 2-cm length; Thermo Fischer Scientific). Peptides were then separated on a C18 column (2-mm particle size, 75-mm inner diameter, 25-cm length; Thermo Fisher Scientific) with a 3-hour gradient starting from 99% solvent A (0.1% formic acid) and ending with 55% solvent B (80% acetonitrile, 0.085% formic acid). The mass spectrometer acquired data throughout the elution process and operated in a data-dependent scheme with full MS scans acquired with the Orbitrap, followed by as many MS/MS ion trap HCD spectra 3 seconds can fit (data-dependent acquisition with top speed mode: 3-s cycle) using the following settings for full MS: automatic gain control (AGC) target value: $1.10e^6$, maximum ion injection time (MIIT): 60 ms, resolution: 6.10e4, m/z range 350–1500. For HCD MS/MS: Quadrupole filtering, Normalised Collision Energy: 30. Ion trap rapid detection: isolation width: 1.6 Th, minimum signal threshold: 5000, AGC: $1.10e^5$, MIIT: 60 ms, resolution: $3.10e^4$. A dynamic exclusion time was set at 30 s. Peptides with charge state less than 1 or greater than 7 were excluded from fragmentation.

Analysis of nLC-MS/MS data: Identifications and quantifications were performed using MaxQuant version 2.0.3.0 (5), the reviewed Human Uniprot database (released March 2020), and a list of frequent contaminant sequences. The false discovery rate was kept below 1% on both peptides and proteins, and a maximum 2 missed cleavages was allowed. Carbamidomethylation of cysteines was set as a constant modification and acetylation of the protein N-terminus and oxidation of methionine were set as variable modifications. Label-free protein quantification (LFQ) was performed using both unique and razor peptides with at least 2 ratio counts. The “match between runs” (MBR) option was allowed with a match time window of 0.7 min and an alignment time window of 20 min. Absolute quantification of cell proteins for intact RBC proteomes was performed using calculated MCH values, as described (6). For RBC ghost proteome absolute quantifications, calculated BAND3 in the corresponding proteome was used as a reference.

Statistical analysis: Data were imported into the Perseus software version 1.6.15.0 (7). Reverse and contaminant and only identified by site proteins were excluded from analysis. Only proteins quantified in at least 3 samples of one condition were selected for two samples Student's T-Test.

**Metabolomic and redox-proteomics**

Metabolomics and redox-proteomics from intact RBCs were performed by ultra-high-pressure liquid chromatography coupled to tandem mass spectrometry (UHPLC-MS/MS).

Sample processing and metabolite extraction: Flow-sorted RBCs were extracted at a concentration of 4 million cells per ml in methanol:acetonitrile:water (5:3:2, *v/v/v*). After vortexing at 4°C for 30 min, soluble extracts were separated from the protein pellet by centrifugation for 10 min at 18,000g at 4°C, and stored at −80°C until analysis.

UHPLC-MS/MS metabolomics: Analyses were performed using a Vanquish UHPLC coupled online to a Q Exactive mass spectrometer (Thermo Fisher, Bremen, Germany). Samples were analyzed using a 1 minute and 5 minute gradients, as described (8–10). Solvents were supplemented with 0.1% formic acid for positive mode runs and 1 mM ammonium acetate for negative mode runs. MS acquisition, data analysis, and elaboration were performed, as described.(8–10)

UHPLC-MS/MS redox-proteomics: Proteomics analyses were performed via Filter Aided Sample Preparation digestion and nano-HPLC-MS/MS identification (TIMS TOF Pro 2 Single Cell Proteomics, Bruker Daltonics, Bremen, Germany), as described (11).

Statistical analyses: Graphs and statistical analyses were prepared with MetaboAnalyst 5.0 (12).

Acronyms in Figure 2 : ACSL : acyl-CoA synthetase; acyl-CX:Y : acylcarnitine with X:Y acyl chain; CMP-Neu5Ac : CMP-N-acetylneuraminate; CoA : Coenzyme A; CPT : carnitine palmitoyltransferase; Cys-Cys : cystine, two cysteines bonded together; Cys-Gly : cysteinyl-glycine; DHAP : dihydroxyacetone phosphate; FA : fatty acid; G6PD : glucose-6-phosphate dehydrogenase; GADP : glyceraldehyde 3-phosphate; Glc-6P : glucose 6-phosphate; Glycerol 3P : glycerol 3-phosphate; GSH/GSSG : reduced/oxidized glutathione; GSR : glutathione-disulfide reductase; LPA/PA : lyso/phosphatidic acyl; LPAT : lysophospholipid acyltransferases; NAD+ : oxidized nicotinamide adenine dinucleotide; NADPH/NADP+ : reduced/oxidized nicotinamide adenine dinucleotide phosphate; Pentose P : Pentose Phosphate, alpha-D-Ribose 1-phosphate was detected here; PL : phospholipid; PLA2 : phospholipaseA2; PPP : pentose phosphate pathway; THcHDO : 3D-(3-5/4)-Trihydroxycyclohexane-1-2-dione.

**Osmotic fragility**

RBC osmotic fragility was determined, as described (2), with modifications to increase sensitivity. Briefly, 0.8 million RBCs were washed in PBS, incubated for 45 minutes in hypotonic NaCl-PO4 solution (equivalent to NaCl solution, ranging from 0% to 0.9%), and centrifuged (800g, 5min). Heme-mediated (non-HRP) peroxidase activity of the released hemoglobin was revealed by adding 50µL of 3,3’, 5,5”-tetramethylbenzidine (TMB) to 12.5µL of supernatant for 1 hour. Absorbance was measured at 655nm using a spectrophotometer, and the percent hemolysis for each salt concentration was calculated.

**Dynamic RBC adhesion on endothelial cells**

Human microvascular endothelial cell line 1 (HMEC-1) cells were seeded at $10^8$ cells/mL in Vena8 Endothelial+ Biochips (Cellix Ltd, Dublin, Ireland), previously coated with 40µL of 0.2% gelatin in PBS. Cells were then incubated for 2 h at 37°C, permitting cell attachment, and then cultured for 48 h using a Kima pump (Cellix Ltd.). RBCs were washed in PBS and resuspended to a 1% Hematocrit in Hank's buffer supplemented with 0.4% bovine albumin, $Ca^{2+}$, $Mg^{2+}$, and HEPES (1mM). To initiate adhesion, a first perfusion step was performed (10 minutes, 0.2 dyn/cm2), enabling interactions between RBCs and endothelial cells. Then the shear stress was increased every 5 minutes (0.5 and then 1 dyn/cm2) to remove less adherent RBCs. Brightfield imaging of adherent RBCs was performed at 10x magnification (AxioObserver Z1, Zeiss). The number of adherent RBCs was determined from 10 pictures taken at the end of the 1 dyn/cm² step for each condition.

**Proteasome activity**

Proteasome activity was measured using Cell-Based Proteasome-Glo™ Assays (Promega), according to the manufacturer's recommendations. Briefly, 100,000 RBCs were mixed with the specific substrate for each activity (Suc-LLVY, Z-LRR, and Z-nLPnLD for chymotrypsin-like, trypsin-like, and caspase-like activity, respectively), diluted in the detection reagent, and incubated for 10 minutes at RT. Luminescence was read with an Infinite 200 Pro (Tecan).

### Supplementary Tables

| Family | | Gene Names | Peak area SS CFSE^low | Peak area LS CFSE^low | Peak area LS CFSE^high |
|---|---|---|---|---|---|
| Proteostasis (24/83 = 29%) | Proteasome | PSMD2 | 2.86 | 5.86 | 5.43 |
| | | PSMD6 | 0.14 | 1.57 | 1.14 |
| | | PSMD13 | 0.29 | 2.57 | 2.00 |
| | | FBXO7 | 0.29 | 3.29 | 3.29 |
| | | CAND1 | 0.57 | 2.43 | 3.14 |
| | | UBE2V1 | 9.71 | 11.57 | 17.00 |
| | | UBE2V2 | 1.43 | 4.00 | 6.43 |
| | | PSMC1 | 0.71 | 3.43 | 2.14 |
| | | PSMD1 | 6.00 | 10.43 | 5.43 |
| | | PSMC2 | 0.00 | 1.57 | 0.43 |
| | | PSMC5 | 0.14 | 1.57 | 0.14 |
| | | PSMA5 | 1.71 | 2.71 | 0.86 |
| | | PSMB7 | 0.14 | 1.57 | 0.57 |
| | | COPS6 | 1.71 | 4.00 | 1.57 |
| | | USP9X | 0.00 | 0.71 | 0.14 |
| | | RNF123 | 1.71 | 4.71 | 3.14 |
| | Chaperones | HSPA8 | 29.43 | 43.71 | 38.00 |
| | | CCT6A | 2.43 | 5.14 | 4.57 |
| | | STIP1 | 5.71 | 10.43 | 13.43 |
| | | HSP90AB1 | 0.00 | 0.43 | 1.14 |
| | | NAP1L4 | 0.43 | 1.00 | 1.14 |
| | | ST13;ST13P5;S | 3.14 | 8.29 | 5.43 |
| | | CCT2 | 10.43 | 19.43 | 13.71 |
| | | CCT4 | 4.43 | 10.14 | 4.57 |
| Cytoskeleton (9/83 = 11%) | | ACTB | 41.57 | 51.00 | 54.71 |
| | | SPTA1 | 107.29 | 134.00 | 128.43 |
| | | SPTB | 129.57 | 154.43 | 162.29 |
| | | ANK1 | 60.43 | 77.00 | 76.00 |
| | | MSN | 0.86 | 4.00 | 2.00 |
| | | JUP | 0.00 | 0.00 | 1.29 |
| | | WDR1 | 2.86 | 2.57 | 4.29 |
| | | CFL1 | 0.00 | 0.14 | 1.43 |
| | | EPB41 | 0.00 | 0.14 | 1.29 |
| Anti-oxidant (7/83 = 8%) | | PRDX1 | 3.29 | 5.71 | 8.86 |
| | | PRDX6 | 12.57 | 20.29 | 12.71 |
| | | BLVRB | 66.14 | 89.86 | 81.14 |
| | | TXNL1 | 0.14 | 1.29 | 0.57 |
| | | CAT | 0.00 | 0.14 | 2.00 |
| | | PARK7 | 0.00 | 0.14 | 1.00 |
| | | PRDX2 | 0.71 | 3.71 | 3.43 |

| Family | Gene Names | Peak area SS CFSE^low | Peak area LS CFSE^low | Peak area LS CFSE^high |
|---|---|---|---|---|
| Glycolysis (5/83 = 6%) | ENO1 | 5.00 | 9.00 | 9.86 |
| | TALDO1 | 4.71 | 6.71 | 5.14 |
| | PGK1 | 25.14 | 28.29 | 32.00 |
| | PFKM | 2.00 | 3.71 | 5.57 |
| | GAPDH | 0.29 | 2.57 | 3.71 |
| Transport (5/83 = 6%) | AGO2 | 1.71 | 3.57 | 2.71 |
| | IPO9 | 0.71 | 2.29 | 0.71 |
| | AP2A1 | 0.00 | 1.57 | 0.71 |
| | AP2B1 | 0.43 | 1.71 | 0.57 |
| | AP2M1 | 0.00 | 0.86 | 0.14 |
| Hemoglobins (3/83 = 3.6%) | HBA1 | 733.00 | 824.14 | 886.00 |
| | HBB | 266.29 | 368.43 | 355.71 |
| | HBD | 32.29 | 39.86 | 53.14 |
| Other (30/83 = 36%) | ACLY | 5.57 | 11.86 | 13.71 |
| | ATP2B4 | 0.00 | 0.71 | 0.86 |
| | FABP5 | 0.29 | 1.43 | 2.43 |
| | GDI2 | 12.86 | 19.86 | 20.14 |
| | EIF4A1 | 9.00 | 12.29 | 12.43 |
| | NME1 | 30.43 | 37.14 | 38.43 |
| | NIF3L1 | 0.29 | 1.57 | 1.57 |
| | PAICS | 2.14 | 5.14 | 4.71 |
| | STOM | 5.86 | 10.43 | 9.57 |
| | ARG1 | 0.00 | 0.14 | 1.00 |
| | CA1 | 55.00 | 71.29 | 90.29 |
| | DNPEP | 0.57 | 0.29 | 1.29 |
| | KRT14 | 8.29 | 12.57 | 19.00 |
| | KRT84 | 0.00 | 0.00 | 1.29 |
| | TSTA3 | 4.00 | 5.29 | 6.43 |
| | TPM3 | 2.29 | 3.14 | 4.43 |
| | KRT1 | 27.00 | 31.57 | 37.57 |
| | ATP6V1A | 2.00 | 4.43 | 2.57 |
| | AARS | 0.29 | 1.86 | 0.57 |
| | THOP1 | 0.00 | 1.71 | 0.00 |
| | SEC14L2 | 0.71 | 2.14 | 1.14 |
| | PNP | 22.43 | 37.00 | 25.86 |
| | LTA4H | 0.00 | 2.43 | 1.57 |
| | MCTS1 | 0.29 | 1.71 | 0.29 |
| | BSG | 0.29 | 1.43 | 0.57 |
| | ADK | 0.00 | 1.86 | 0.43 |
| | ANXA7 | 3.86 | 9.29 | 4.86 |
| | YWHAB | 2.86 | 5.00 | 3.00 |
| | YWHAG | 3.86 | 6.00 | 1.43 |
| | CD44 | 0.86 | 1.14 | 0.00 |

**Supplementary Table 1: Classification of oxidized proteins during storage.** Gene names from the reviewed Human Uniprot database are indicated. Peak areas are indicated for each detected oxidized protein. Proteins denoted in black are reversibly oxidized, whereas those denoted in brown are irreversibly oxidized (Cys to DHA).

### Supplementary Figures

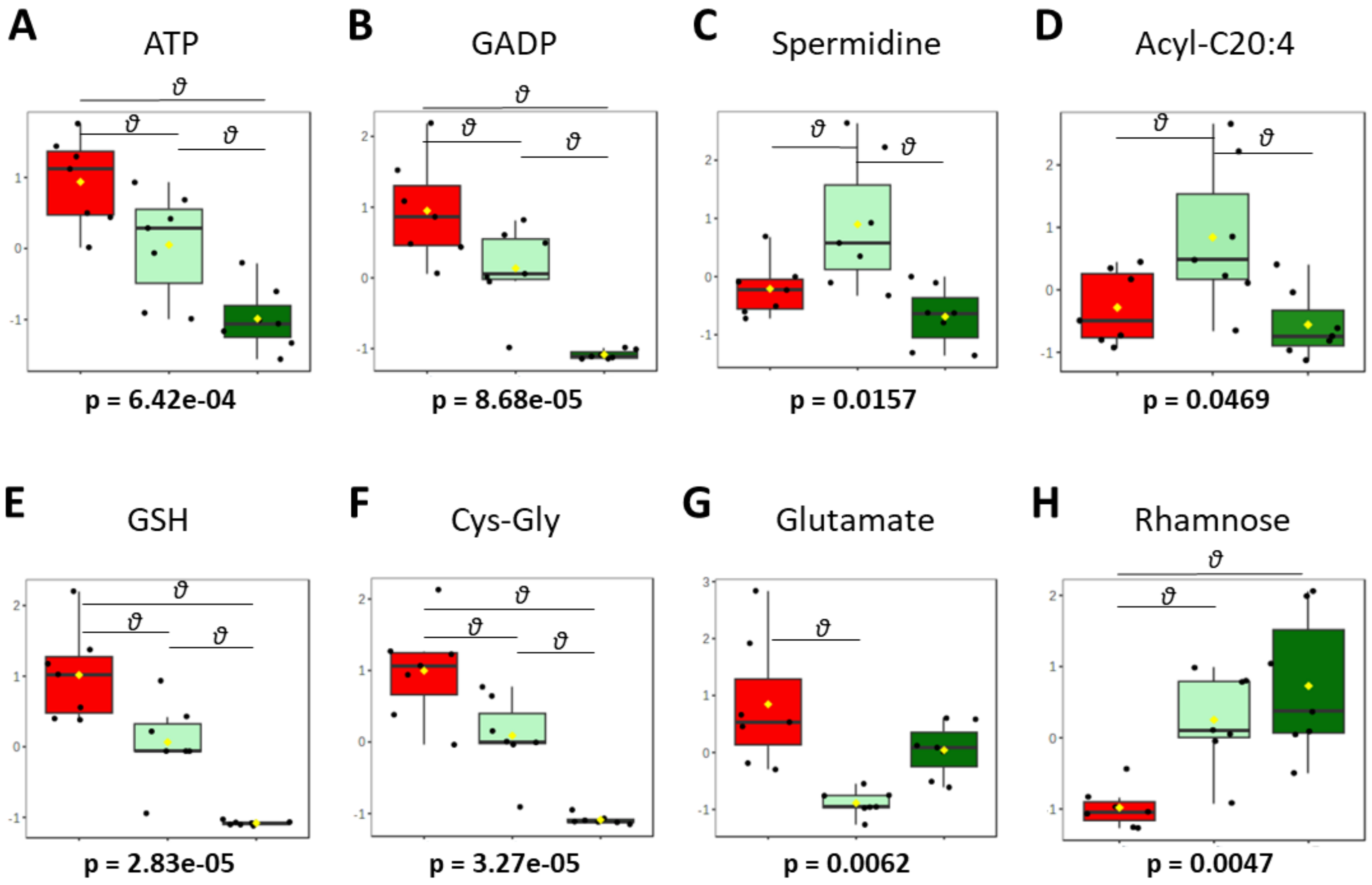


**Supplementary Figure 1: Metabolites that significantly vary between flow-sorted short-stored CFSE$^{low}$ and long-stored CFSE$^{low}$ RBCs.** Box plots from metabolomic data for metabolites that significantly vary between flow-sorted short stored-CFSE$^{low}$ (Red box plots; n = 7) and long stored-CFSE$^{low}$ (Light green box plots; n = 7) RBCs and for flow-sorted long-stored CFSE$^{high}$ RBCs (Dark green box plots; n = 7). P-values of ANOVA are written under each graph and $\theta$ represents a significant difference found by a positive post-hoc test of Tukey's HSD between groups. GADP : glyceraldehyde 3-phosphate; Acyl-C20:4 : acylcarnitine with a 20:4 acyl chain; GSH : reduced glutathione; Cys-Gly : cysteinyl-glycine

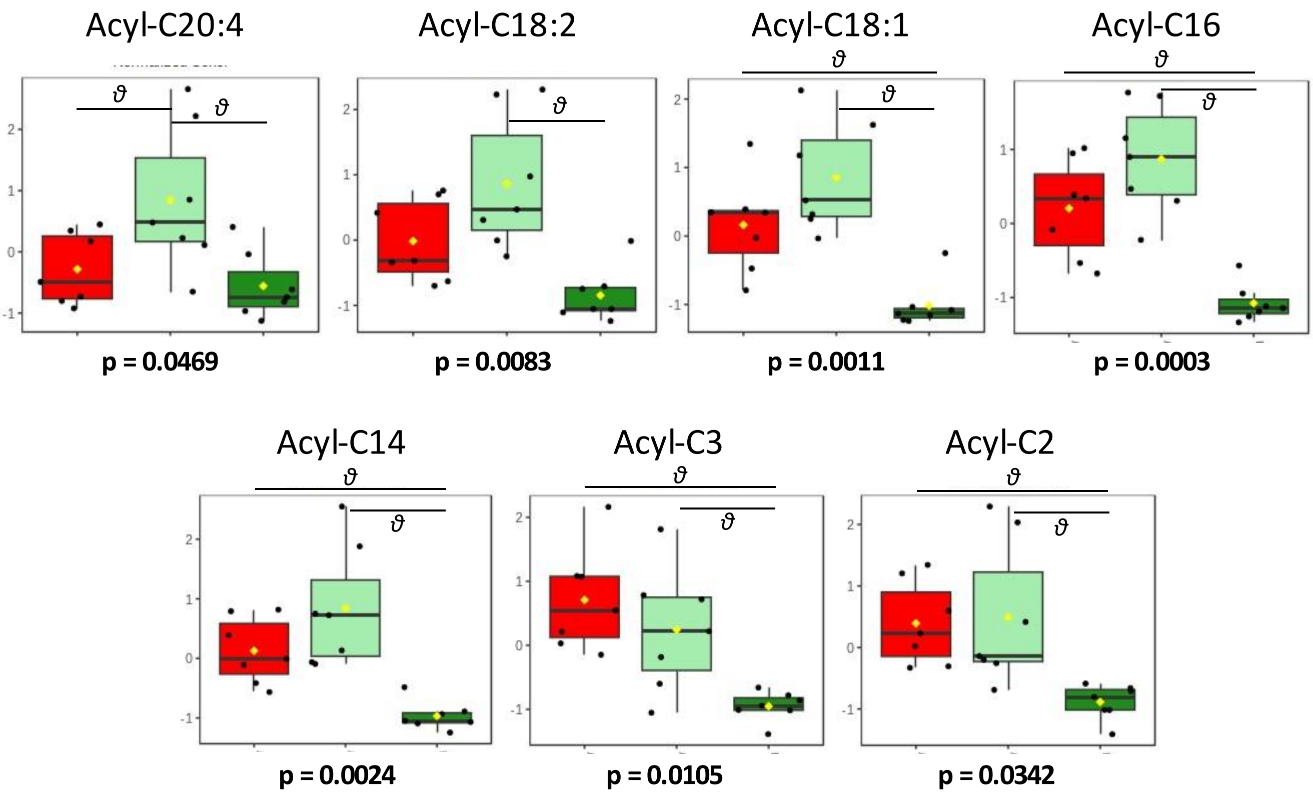


**Supplementary Figure 2: Acyl-carnitines that significantly vary between flow-sorted CFSE-stained RBC subsets.** Box plots of metabolomic data for metabolites that significantly vary between flow-sorted short stored-CFSE$^{low}$ (Red box plots; n = 7), long stored-CFSE$^{low}$ (Light green box plots; n = 7), and long-stored CFSE$^{high}$ RBCs (Dark green box plots; n = 7). P-values of ANOVA are written under each graph and $\theta$ represents a significant difference found by a positive post-hoc test of Tukey's HSD between groups. Acyl-CX:Y means acylcarnitine with a X:Y acyl chain

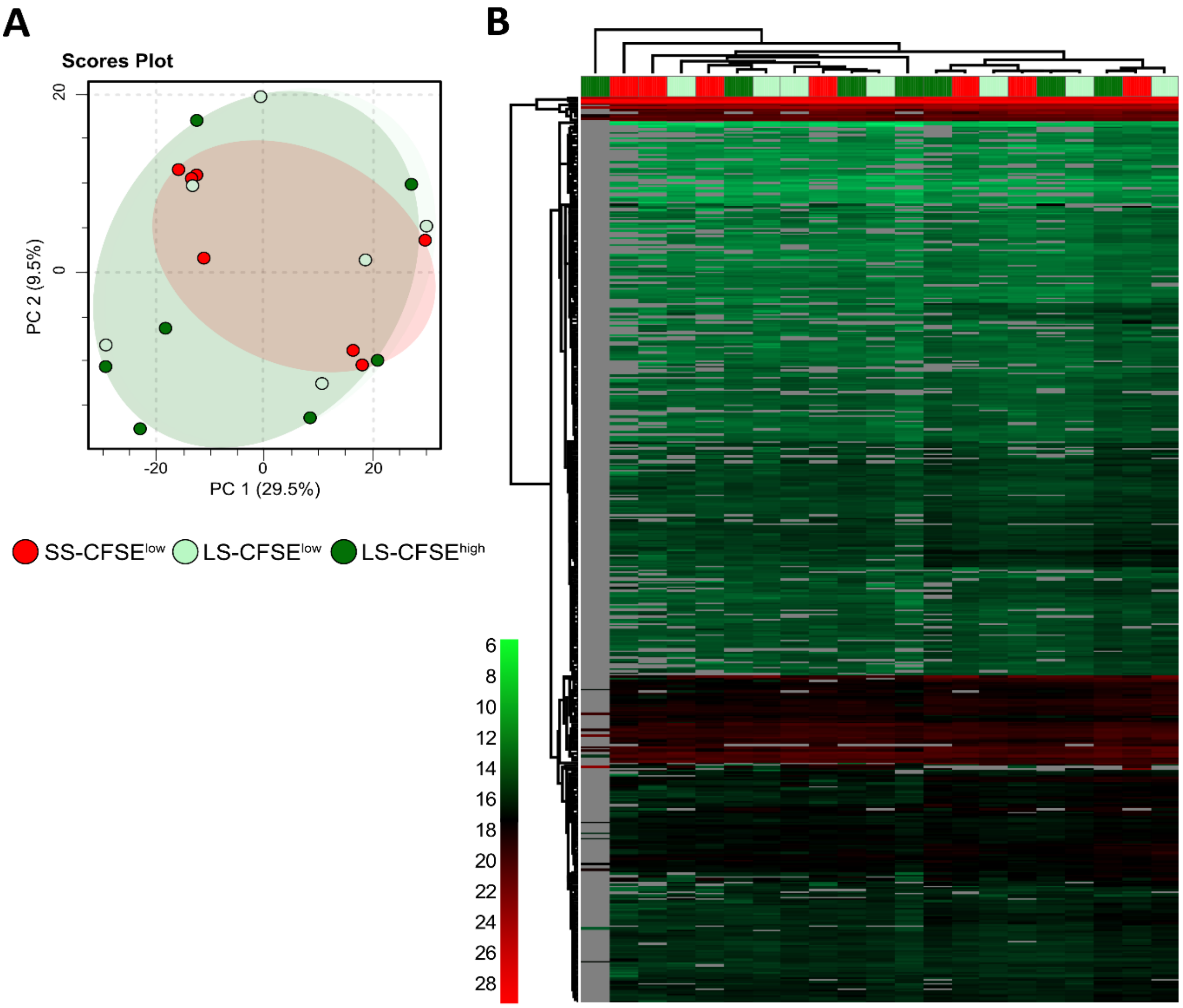


**Supplementary Figure 3: Proteomics of intact RBCs reveal no major differences between the 3 CFSE-stained, flow-sorted, RBC subsets analyzed. (A)** Principal component analysis of proteomics data for flow-sorted short-stored CFSE$^{low}$ (red, n = 6), long-stored CFSE$^{low}$ (light green, n = 6), and long-stored CFSE$^{high}$ RBCs (dark green, n = 6). **(B)** Hierarchical clustering analysis of the proteins (quantified in at least 70% of the samples in one condition) shows no clustering of the three subsets (data are represented by copy number per cell).

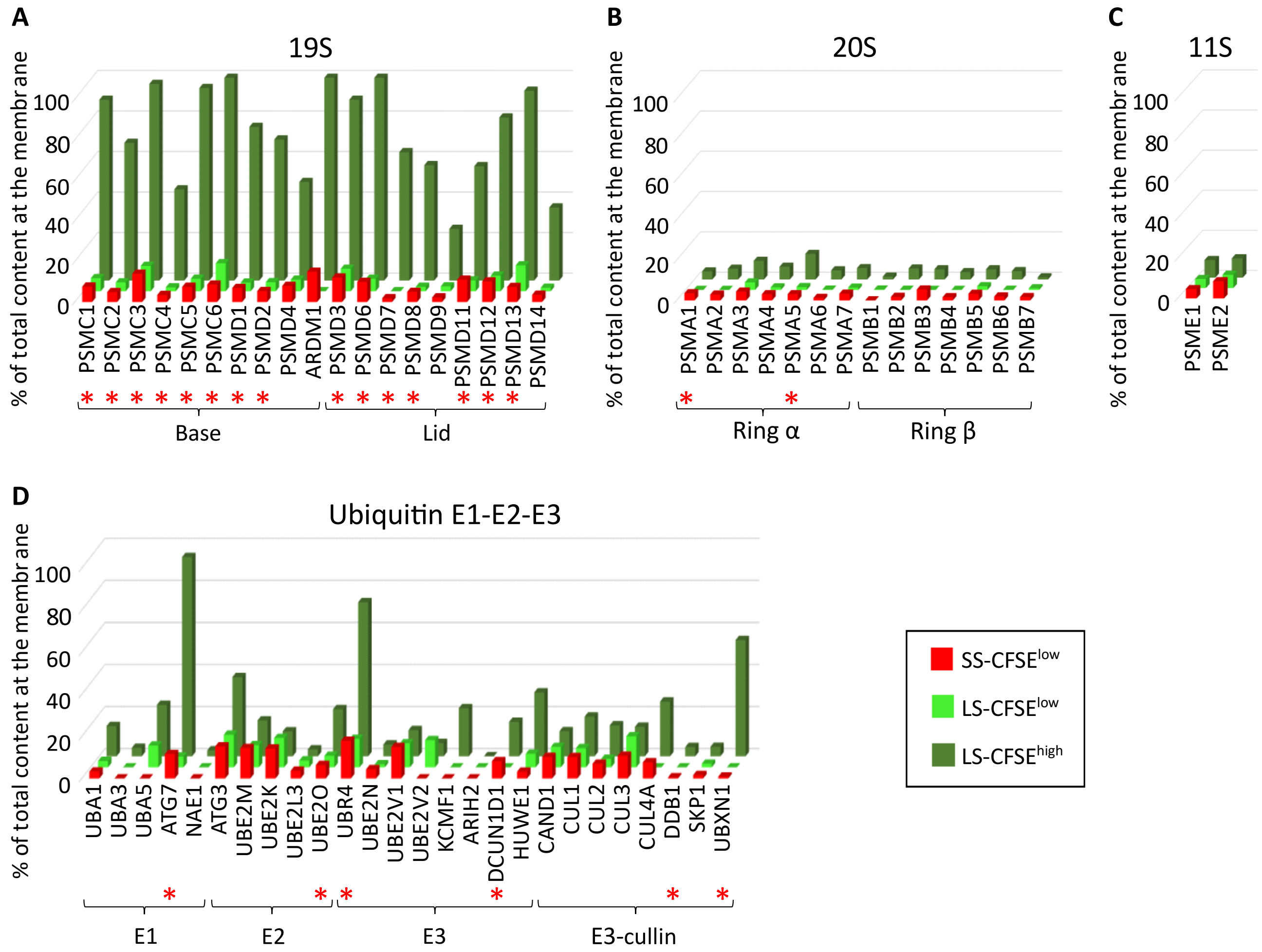


**Supplementary Figure 4: Membrane relocation of RBC proteasome and ubiquitin proteins.** The proportions of total cellular content found at the membrane of flow-sorted short-stored CFSE$^{low}$ (SS-CFSE$^{low}$, in red), long-stored CFSE$^{low}$ (LS-CFSE$^{low}$, in light green), and long-stored CFSE$^{high}$ (LS-CFSE$^{high}$, in dark green) RBCs for all detected proteins in the 19S **(A)**, 20S **(B)**, and 11S **(C)** proteasome subunits, and the E1, E2, and E3 ubiquitin enzymes **(D)**. Red stars identify proteins that show statistically significant membrane relocation in long-stored CFSE$^{high}$ RBCs (*vs* long-stored CFSE$^{low}$ RBCs).

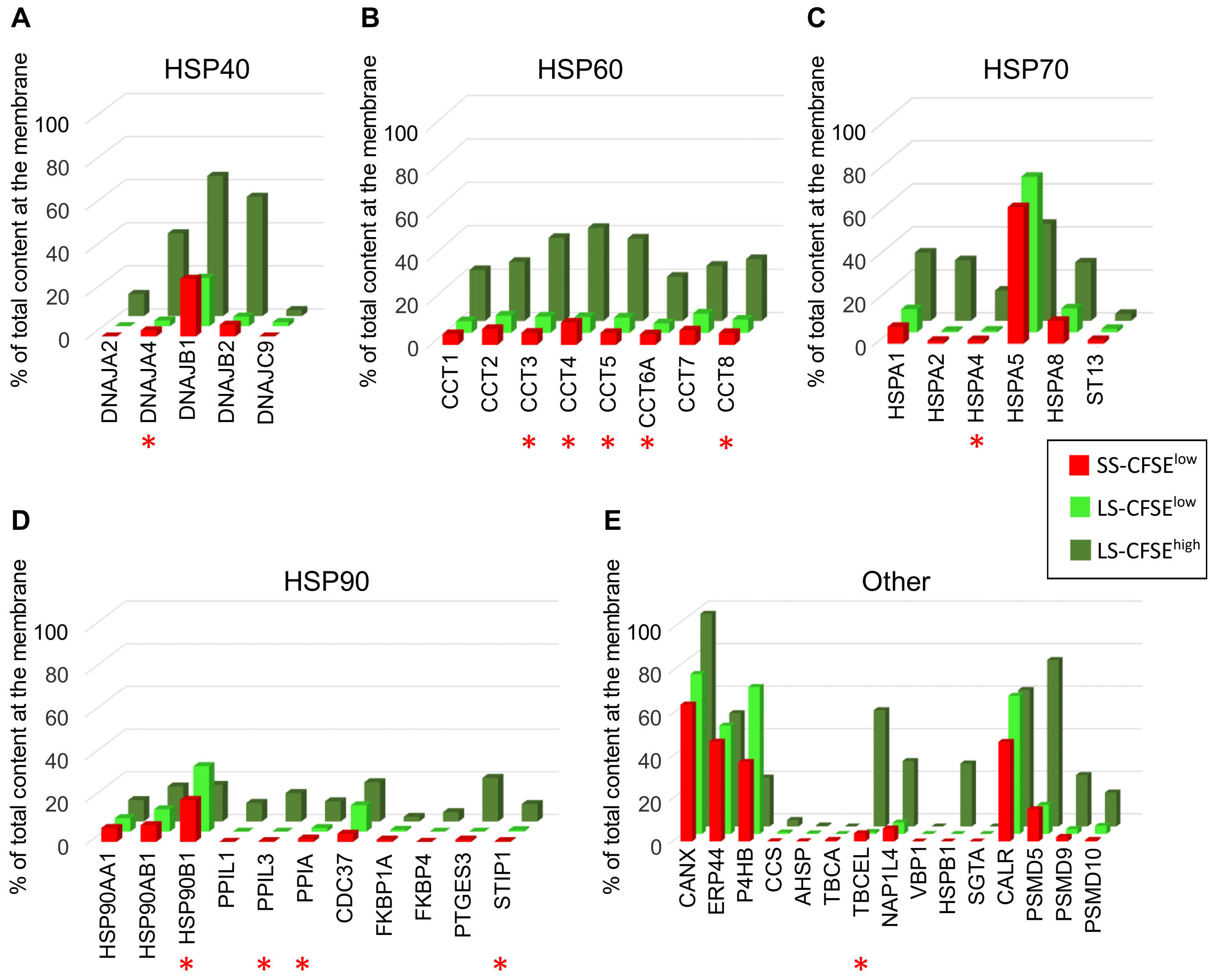


**Supplementary Figure 5: Membrane relocation of RBC chaperone proteins.** The proportions of the cellular total content found at the membrane of flow-sorted short-stored CFSE$^{low}$ (SS-CFSE$^{low}$, in red), long-stored CFSE$^{low}$ (LS-CFSE$^{low}$, in light green), and long-stored CFSE$^{high}$ (LS-CFSE$^{high}$, in dark green) RBCs for all detected chaperone proteins in the **(A)** HSP40, **(B)** HSP60, (**C**) HSP70, and (**D**) HSP90 families, and (**E**) other chaperone proteins. Red stars identify proteins that show statistically significant membrane relocation in long-stored CFSE$^{high}$ RBCs (*vs* long-stored CFSE$^{low}$ RBCs).

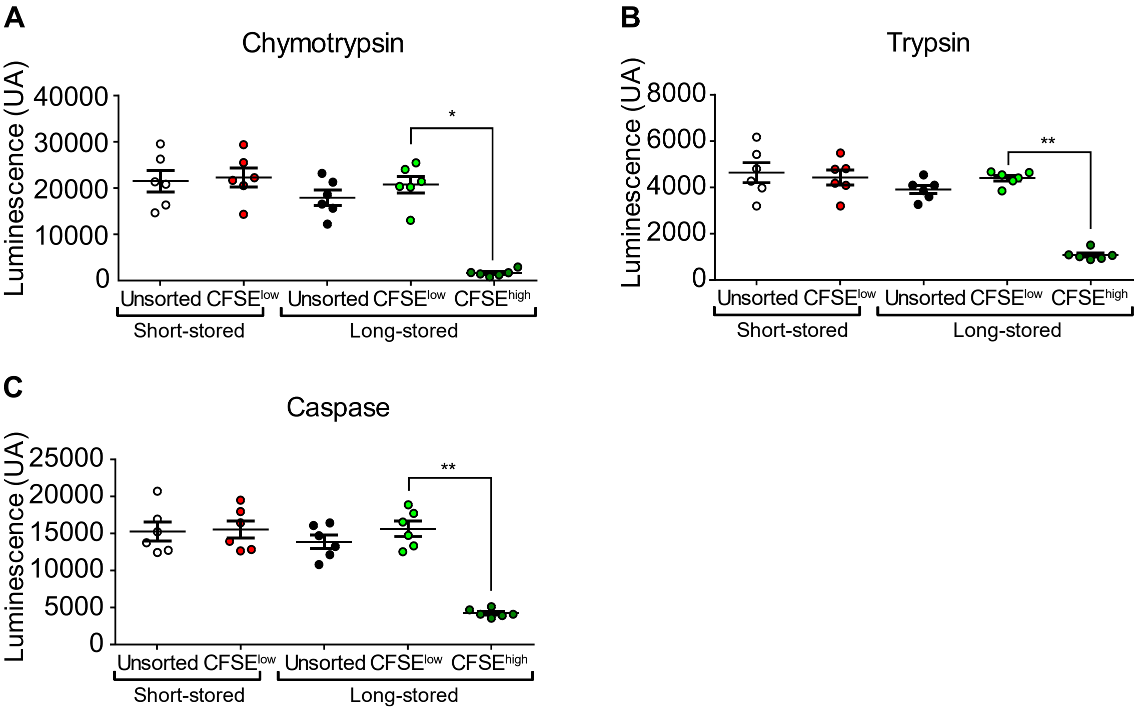


**Supplementary Figure 6: Short-stored CFSE$^{low}$ and long-stored CFSE$^{low}$ RBCs have similar proteasomal activity.** Chymotrypsin-like (**A**), Trypsin-like (**B**), and Caspase-like (**C**) proteasome activities were measured for CFSE-stained short-stored (unsorted and CFSE$^{low}$) and long-stored (unsorted, CFSE$^{low}$, CFSE$^{high}$) RBC subsets. Data are presented as 6 individual experiments with the mean shown ± SEM. * P <0.05, ** P <0.01 by Friedman one-way ANOVA followed by Dunn's multiple comparison test.

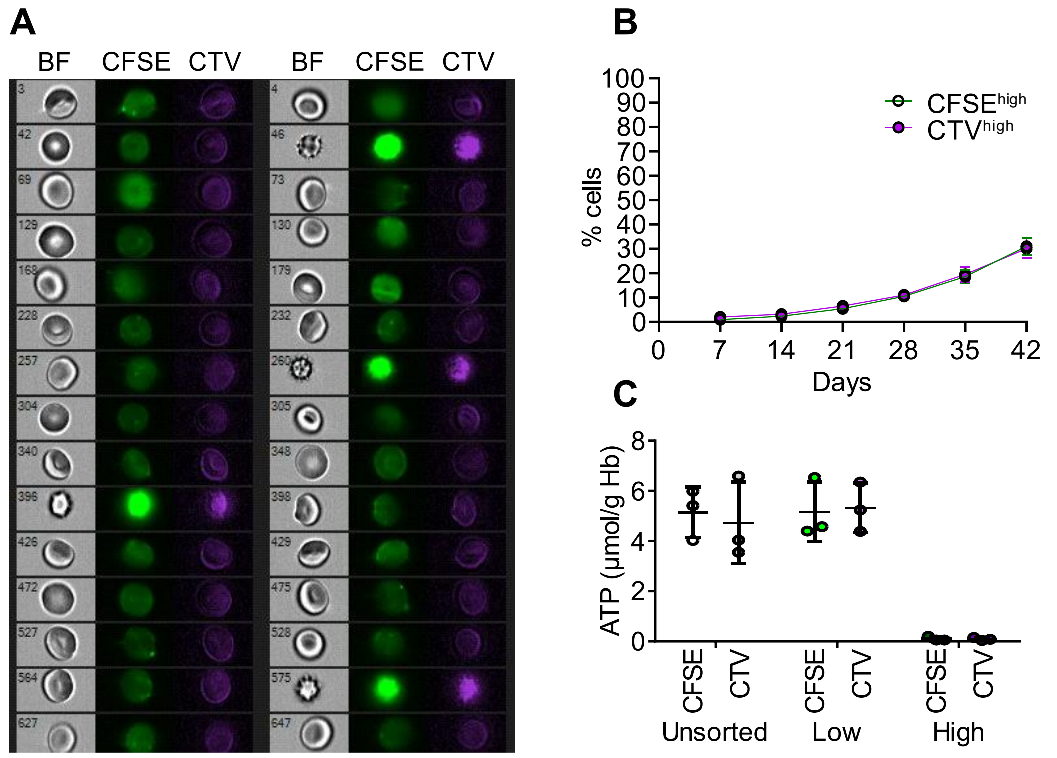


**Supplementary Figure 7: The CTV probe shows staining properties similar to CFSE, allowing quantification of morphologically-altered RBCs by flow cytometry in a different fluorescence channel. (A)** Representative images of CFSE and CTV co-stained RBCs obtained by ImageStream; BF = Brightfield. **(B)** Quantification of CFSE$^{high}$ and CTV$^{high}$ RBCs cells upon storage of RBC concentrates in SAGM solution for 42 days (n = 8, mean ± SEM). **(C)** Intracellular ATP levels normalized for hemoglobin content in sorted RBC subsets identified by CFSE or CTV staining. Data are presented as 3 individual experiments with the mean shown ± SD. In panels B and C, a two-way ANOVA followed by a Sidak's multiple comparison was performed to compare both staining at each time points or for each subset, respectively; no statistically significant differences were observed.

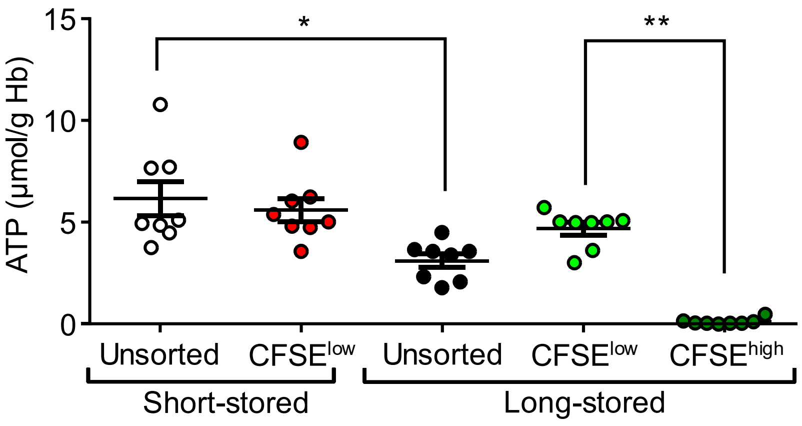


**Supplementary Figure 8: Flow-sorted short-stored CFSE$^{low}$ RBCs and long-stored CFSE$^{low}$ RBCs have similar intracellular ATP levels.** Intracellular ATP levels normalized for hemoglobin content were evaluated for CFSE-stained short-stored (unsorted and CFSE$^{low)}$ and long-stored (unsorted, CFSE$^{low}$, CFSE$^{high}$) RBC subsets. Data are represented as 8 individual experiments with the mean shown ± SEM. * P <0.05, ** P <0.01 by Friedman one-way ANOVA followed by Dunn's multiple comparison test.
